# Translating Histopathology Foundation Model Embeddings into Cellular and Molecular Features for Clinical Studies

**DOI:** 10.64898/2026.03.17.711896

**Authors:** Saishi Cui, Zhining Sui, Ziyi Li, Kristina A. Matkowskyj, Ming Yu, William M. Grady, Wei Sun

## Abstract

AI-powered pathology foundation models provide general-purpose representations of histopathological images by encoding image tiles into numerical embeddings. However, these embeddings are not directly interpretable in biological or clinical terms and must be translated into biologically meaningful features, such as cell-type composition or gene expression, to enable downstream clinical applications. To bridge this gap, we developed STpath, a framework that integrates histopathology image embeddings derived from existing pathology foundation models with matched, spatially resolved transcriptomics data. STpath consists of cancer-specific XGBoost models trained to infer cell-type compositions and gene expression from histopathology image tiles. We evaluated STpath in colorectal and breast cancer datasets and showed that it provides accurate estimates of the composition of major cell types and the expression of a subset of genes, with further performance gains achieved by combining embeddings from multiple foundation models. Finally, we demonstrated that STpath inferred features that can be used in downstream studies to evaluate their associations with clinical outcomes.

## Introduction

Histopathology images, most commonly stained with hematoxylin and eosin (H&E), are routinely used for cancer diagnosis, prognosis, and treatment selection. The rapid growth of large-scale digital pathology repositories^1^, together with the proven success of deep learning in medical image analysis, positions digital pathology as one of the biomedical domains most ready for transformation by artificial intelligence (AI). Recently, several foundation models^2–6^ have been developed and trained on large collections of H&E images. However, by design, these models are general-purpose and generate numerical embeddings that are often difficult to interpret, limiting their utility for specialized tasks such as dissecting the tumor microenvironment. As pathology foundation models are increasingly used in translational and clinical studies, there is a timely need for approaches that can translate their abstract representations into interpretable features suitable for clinically-oriented studies and ultimately to be used in clinical care.

To bridge this gap, we developed STpath, a computational framework to train STpath models that map numerical embeddings of pathology foundation models to biologically meaningful features, such as cell-type composition or gene expression. Each STpath model operates at the image-tile level, taking an H&E tile as input and predicting corresponding features. At its core, STpath leverages XGBoost models trained on the embedding of H&E images and the matched, spatially resolved transcriptomic (SRT) data, providing a robust and interpretable strategy for translating image representations into cellular or molecular phenotypes. Our work makes three primary contributions. First, we deliver a collection of pre-trained STpath models for colorectal and breast cancers, enabling direct investigation of cell-type composition or gene expression from H&E images in these disease contexts. Second, we build a flexible and user-friendly computational framework that facilitates the training of STpath models for additional cancer types when matched SRT data are available. Third, through a systematic evaluation of multiple pathology foundation models, we demonstrate that integrating embeddings from different models further improves the accuracy of feature prediction from H&E images.

Recent studies^7^ have suggested that embeddings derived from pathology foundation models may capture not only biological signals but also technical or contextual variations, such as tissue site or medical institution. In addition, the image tiles extracted from the same H&E image may be similar to each other due to image-level batch effects. Such batch effects can obscure the inferred features. These challenges highlight the need for computational frameworks that are sufficiently robust to overcome batch effects and enable recovery of biologically meaningful molecular and cellular features from pathology foundation model embeddings.

To address these challenges and to assess the robustness and generalizability of our approach, we systematically evaluated STpath models using embeddings from five representative pathology foundation models: Conch^2^, Prov-GigaPath^3^, UNI2-h^4^, Virchow^5^, and Virchow2^6^. Several prior methods^8–17^ have explored the prediction of gene expression from H&E images using neural network-based approaches, and a recent benchmark study^18^ identified that DeepPT^10^, a ResNet50-based model, performed better than or comparable to other methods. Accordingly, we included ResNet50-based embeddings as a comparator in our analyses.

Our work represents the first systematic assessment of pathology foundation models and their combinations in their ability to resolve cellular neighborhoods within the tumor microenvironment. Such evaluation is essential for the responsible translational use of foundation models, as unrecognized technical artifacts can confound biological interpretation and ultimately influence clinical decision-making. By explicitly characterizing the susceptibility of existing foundation models to batch effects and introducing principled strategies to mitigate them, our study establishes a foundation for more reliable inference from histopathology images. Using colorectal and breast cancer H&E datasets, we demonstrate that STpath offers a robust and generalizable framework for translating image embeddings into biologically and clinically meaningful features, facilitating large-scale downstream association studies. Finally, we provide a well-documented workflow that can serve as a practical blueprint for adapting this approach to additional cancer types and other molecular/cellular endpoints.

## Results

### Generating High-Quality Labels for Model Training

All H&E foundation models considered in this study divide the whole slide image into a series of image tiles and output embeddings for each tile, typically around 256 x 256 pixels at 20x magnification. Our goal was to develop a machine learning method that converts such tile-level embeddings into cell type compositions (cellular features) or gene expression profiles (molecular features). Since it is not feasible to manually annotate tens of thousands of image tiles, we trained STpath using paired H&E images and SRT data. When cellular- or subcellular-resolution SRT data were available (e.g., 10x Xenium), cell type proportions could be directly calculated for each tile. Currently, most publicly available SRT data are spot-level (e.g., 10x Visium), where one spot may cover 1 to 10+ cells. In such cases, we performed cell type deconvolution using matched single-cell sequencing (scRNA-seq) data as cell-type-specific references. This workflow, including the crucial steps of aligning H&E-stained whole slide images (WSIs) with SRT spot coordinates and generating matched tissue tiles, are schematically outlined in Figure 1A and 1B. In total we have collected 63 H&E images covering more than 120,000 image tiles for colorectal cancer, and 11 H&E images covering more than 43,000 image tiles for breast cancer data (Supplementary Table 1-2).

**Figure 1.**
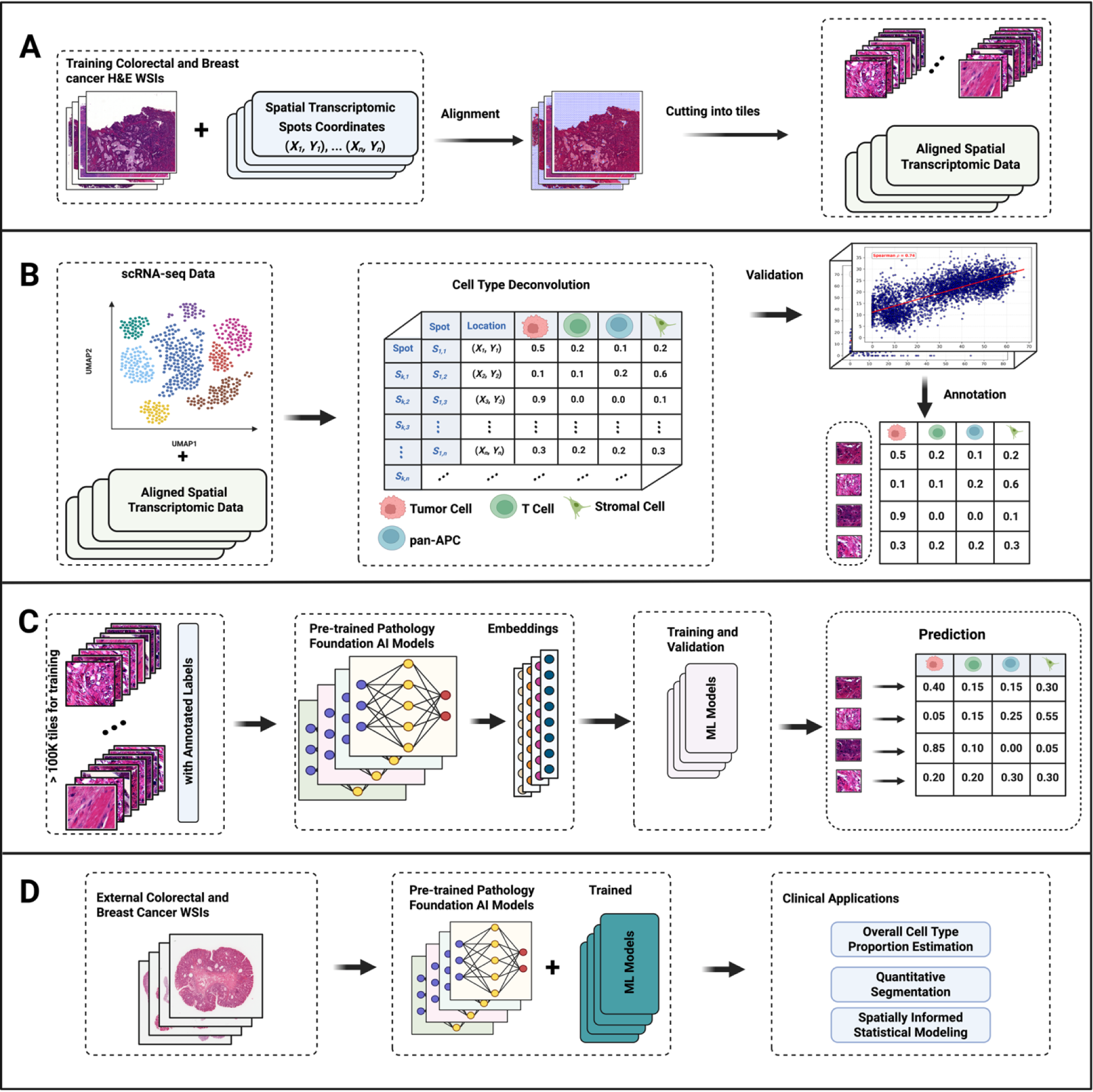
Overview of the STpath. (A) Whole-slide H&E images were aligned with spatial transcriptomics spot coordinates and partitioned into tissue tiles. (B) scRNA-seq data were processed to curate reference cell type signatures, followed by cell type deconvolution of ST data and validation of inferred cell type proportions with marker gene expression. Annotated tiles were generated by linking deconvolution results to matched image regions. (C) Annotated tissue tiles were used to extract embeddings from multiple pre-trained pathology foundation AI models. These embeddings were input into ML models to predict cell type proportions, with training and leave-one-individual-out validation. (D) Trained models were applied to external colorectal and breast cancer WSIs, enabling downstream clinical applications including overall tumor grading, quantitative soft tissue segmentation, and association with genomic and clinical outcomes.

Using a customized cell-type-specific gene expression reference (see Methods Section for details), we performed cell type deconvolution on the SRT data with CARD^21^. For colorectal cancer, this analysis yielded deconvoluted proportions for 19 cell subtypes (Figure 2C, top), which were then aggregated into five major categories: Tumor Cells, Normal Epithelial Cells, T Cells, pan-antigen-presenting-cells (pan-APCs), and Stromal Cells (Figure 2C, bottom). We separated T cells from other immune cells due to their importance in cancer treatment and prognosis. Since other immune cells are all antigen presenting cells, we refer to them as pan-antigen-presenting-cells or pan-APC. Further separating other immune cells to more refined cell types reduced prediction accuracy, likely due to low proportions of each cell type (Figure S1A). We validated the accuracy of deconvolution by comparing the estimated proportions against the expression of selected cell type-specific marker genes, confirming that the proportion of a given cell type was strongly correlated with the expression of its corresponding marker genes in most images (Figure 2D). Detailed cell type deconvolution results for each sample, including mean cell type proportions, as well as quantitative validation metrics, are provided in Supplementary Files 1–2. We further filtered out images with low correlations between marker gene expression and estimated cell type proportions according to the criteria described in Supplementary Methods: Deconvoluted Results Validation. Briefly, Spearman correlations were computed between deconvoluted cell type proportions and relative marker gene expression, and samples with correlations exceeding the threshold defined for each cell type were retained. The retained images were then manually inspected to confirm that estimated cell type proportions and marker gene expression showed consistent spatial distributions on the H&E images (Figure 2E).

**Figure 2.**
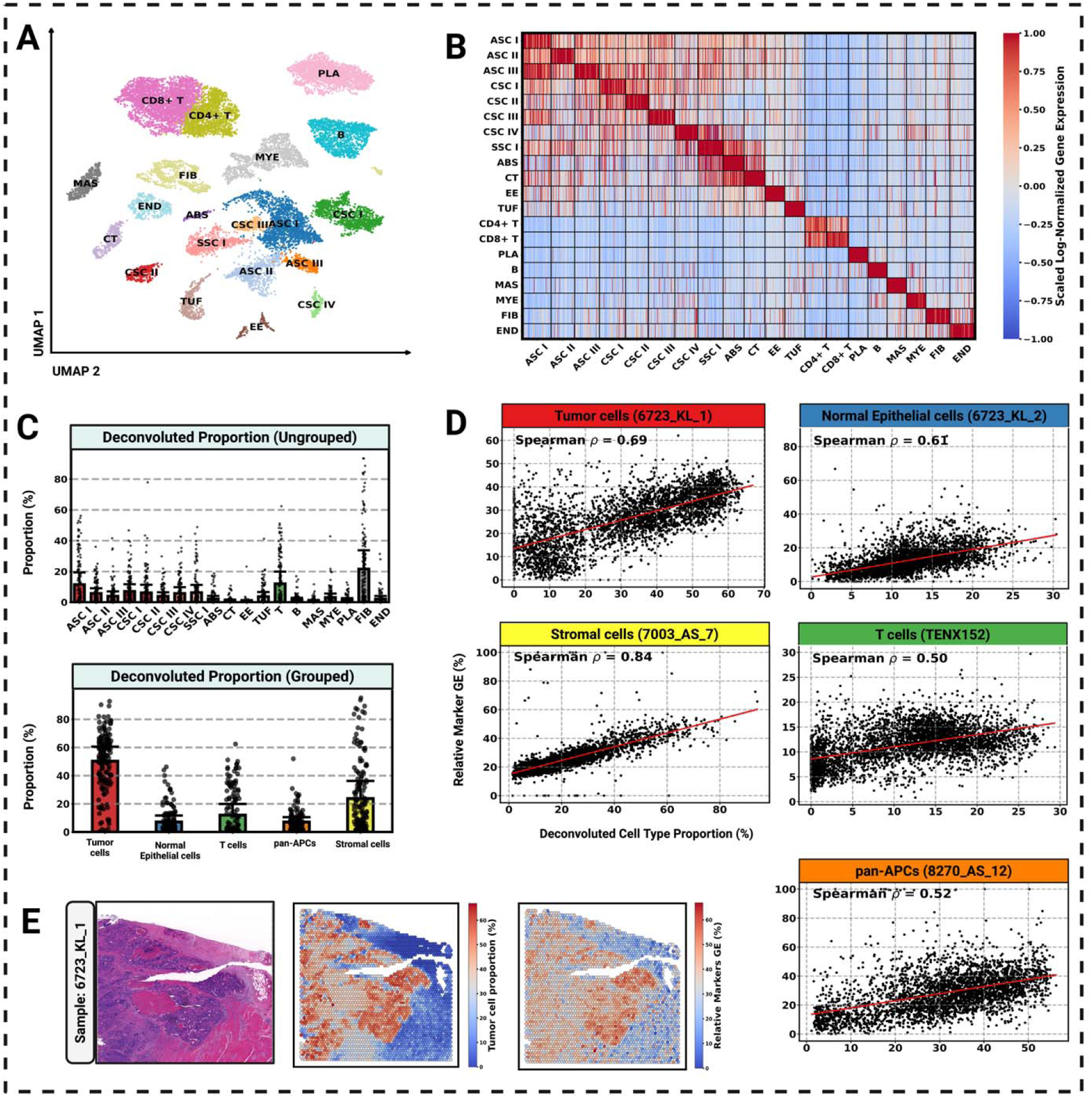
Cell type deconvolution and validation. (A) UMAP visualization of double-confirmed colorectal cancer single-cell RNA-seq data showing 20 annotated cell subtypes, including tumor subtype (ASC: Absorptive stem-like cancer cells, CSC: Carcinoma-specific epithelial cells, SSC: Serrated-lesion–derived cells), normal epithelial subtypes (CT: Crypt-top colonocytes, EE: Enteroendocrine cells, TUF: Tuft cells, ABS: Absorptive colonocytes), stromal subtypes (FIB: Fibroblasts, END: Endothelial cells), T cell subtypes (CD4+T, CD8+T) and pan-APC subsets (PLA: Plasma, MYE: Myeloid, MAS: Mast cells, B cells). (B) Heatmap of scaled log-normalized marker genes expression across cell subtypes, confirming distinct transcriptional profiles. (C) Deconvoluted cell type proportions at the tissue region level, shown for ungrouped subtypes (top) and aggregated into five major categories (bottom). (D) Correlation analysi between deconvoluted cell type proportions and marker gene expression–based relative expression. Representative examples show positive Spearman correlations across all five major categories, supporting the validity of deconvolution results. (E) Representative H&E-stained image with corresponding spatial hexagonal binned heatmaps of deconvoluted tumor cell proportions and relative marker genes expression (GE).

### Supervised Learning Can Mitigate the Batch Effects of Foundation Models

We first extracted high-dimensional numerical embeddings from each tile using five pathology foundation models (Conch, Prov-GigaPath, UNI2-h, Virchow, and Virchow2) as well as ResNet50. We refer to each dimension of the embeddings as a feature. For example, UNI2-h outputs a 1,536-dimensional embedding vector for an image tile and thus it has 1,536 features. To visualize these high-dimensional feature spaces, we applied UMAP to the embeddings from each model, with each point representing a single tile image. When using the full feature sets, UMAP revealed noticeable batch effects, with tiles from the same individual clustering together (Figure 3A). This batch effect was even more pronounced in some foundation models, such as Virchow2, compared to the ResNet50 baseline, likely because pathology foundation models are pretrained on large-scale histopathology datasets and tend to encode rich patient- and slide-specific morphological signatures, which can amplify inter-individual variability in the embedding space.

**Figure 3.**
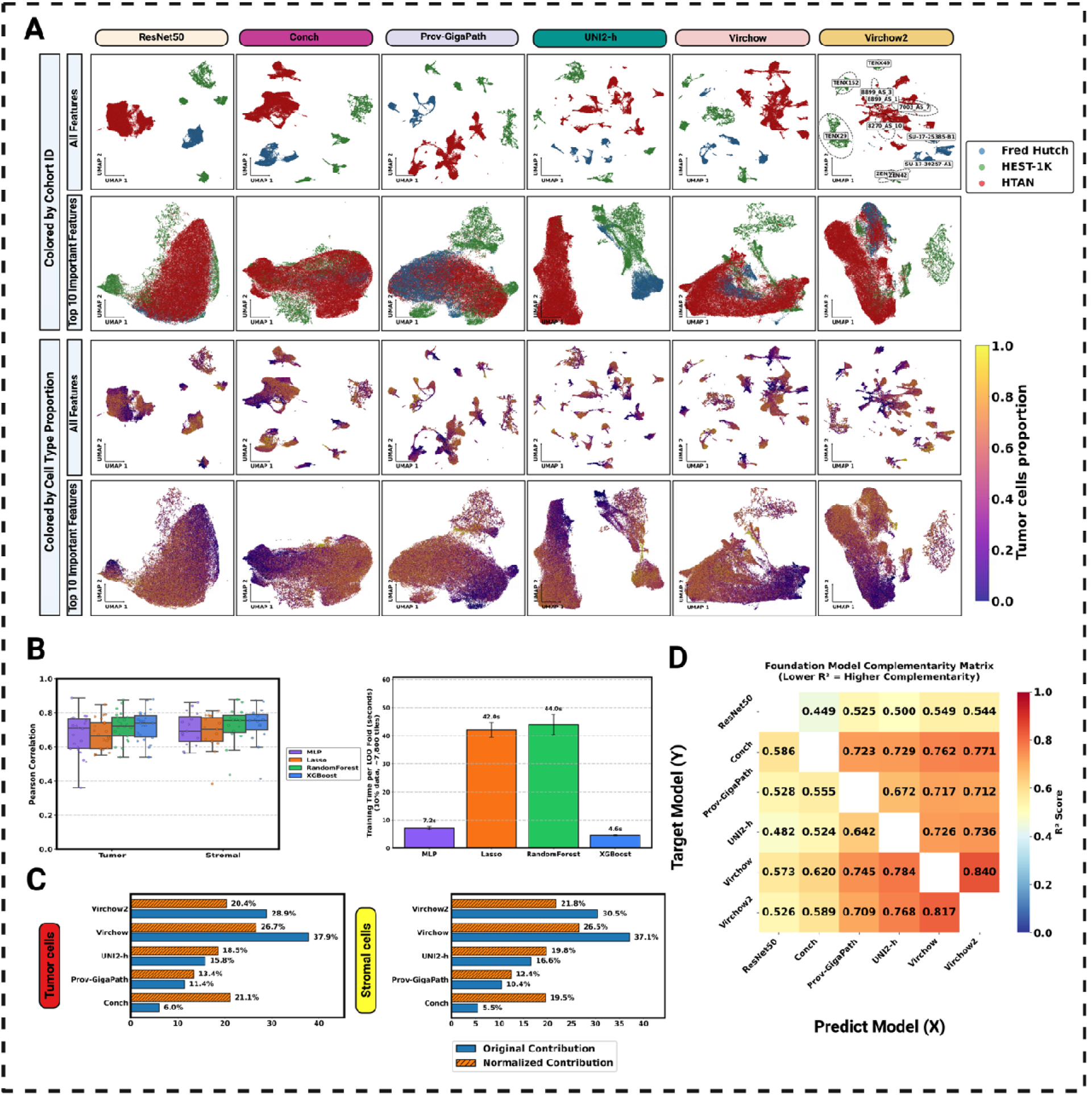
Feature representation, predicting models’ comparison, efficiency, and complementarity across pathology foundation models. (A) UMAP visualization of extracted features from 5 pathology foundation models and ResNet50 as a reference model. For each model, both the full features set, and the top 10 features selected by XGBoost were visualized, with embeddings colored by dataset cohorts (top) and deconvoluted cancer cell proportions (bottom). (B) Contribution analysis of foundation models to the combined XGBoost prediction performance for tumor and stromal cells. Both absolute contributions (blu ) and normalized per-feature contributions (orange) are shown, highlighting complementary strengths of different models. (C) Comparison of predictive performance and computational efficiency across multiple machine learning methods, including MLP, Lasso, Random Forest, and XGBoost. Models were evaluated using leave-one-out training. (D) Pairwise complementarity analysis of foundation model features using ridge regression. The heatmap shows scores between predicted and target models, with lower values indicating higher complementarity.

While this batch effect could be alleviated by aligning them to the same space, similar to the methods for aligning different types of single-cell omics data, we found feature selection by supervised learning is more effective at removing batch effects. We trained separate predictive models for each cell type to estimate cell type proportions from the embedding features of each foundation model. Among the supervised learning methods evaluated, including feed-forward neural networks, random forests, and XGBoost^22^, both random forest and XGBoost achieved consistently strong and comparable performance, outperforming other models (Figure 3C, left). As ensemble-based methods, these models are well-suited for heterogeneous pathology-derived features, offering improved generalization and reduced susceptibility to overfitting under limited training data. Notably, XGBoost demonstrated substantially higher computational efficiency while maintaining similar predictive accuracy to random forests (Figure 3C, right). Therefore, we selected XGBoost for all subsequent analyses to enable systematic comparisons of foundation models across cancer types. Built-in feature importance metric (gain) from the trained XGBoost models was used to identify sets of informative features. When restricting the analysis to the ten most important features selected by XGBoost, the individual-level batch effects were markedly reduced, resulting in tiles being grouped by their cell type proportions (Figure 3A, Figure S2-S3). These results highlight that supervised learning can identify the most relevant features for downstream tasks while mitigating batch effects.

### Different Foundation Models Provide Complementary Information

Because distinct foundation models were trained on different datasets using diverse modeling strategies, we hypothesized that they capture complementary information that could be integrated to improve predictive performance. Towards this end, we created a combined model by concatenating features from all five foundation models. Feature importance analysis of the combined model was then used to assess the relative contributions of each foundation model. We quantified these contributions using two metrics: absolute contribution, which measures the total predictive power of each model; and normalized contribution, which accounts for the varying number of features contributed by each model and thus reflects per-feature efficiency.

Contributions varied across cell types. For instance, Virchow and Virchow2 showed the highest absolute contributions when predicting tumor cell proportions, whereas Prov-GigaPath and UNI2-h had the highest normalized contributions for T-cell proportion prediction (Figure 3B, Figure S1B). These results indicate that individual foundation models capture distinct aspects of cell morphology that are relevant to specific cell types.

To further assess the complementarity of feature representations, we performed pairwise ridge regression to predict the features of one model from those of another. A high *R*^2^ value between two models indicates substantial redundancy in their features. As expected, Virchow and Virchow2, which were developed by the same researcher team as successive versions, showed high *R*^2^ values (0.84/0.82; Figure 3D), confirming their features are less complementary. In contrast, ResNet50, trained on images from ImageNet, showed low *R*^2^ values relative to most pathology foundation models (Figure 3D). Therefore, the pathology-specific foundation models capture unique information beyond those from ResNet50, and that different foundation models contribute complementary signals.

### STpath Gives Accurate Estimation of Cell Type Proportions

We evaluated the trained XGBoost models using a stringent leave-one-individual-out (LOIO) cross-validation scheme to prevent data leakage and to ensure that model performance reflects true generalization across individuals. Model performance was assessed using two complementary metrics: mean absolute error (MAE), with smaller values indicating higher accuracy, and the Pearson correlation coefficient between predicted and deconvoluted proportions, with larger values indicating a stronger linear relationship.

ResNet50 consistently exhibited inferior performance, characterized by higher MAE and lower Pearson correlations, compared with the pathology foundation models (Figure 4A), highlighting the stronger representational capacity of the foundation models pretrained on large histopathology datasets. Among the foundation models, Virchow and Virchow2 achieved relatively better performance. Notably, the combined model, which integrates important features from all foundation models, consistently outperformed any single model across all target cell types.

**Figure 4.**
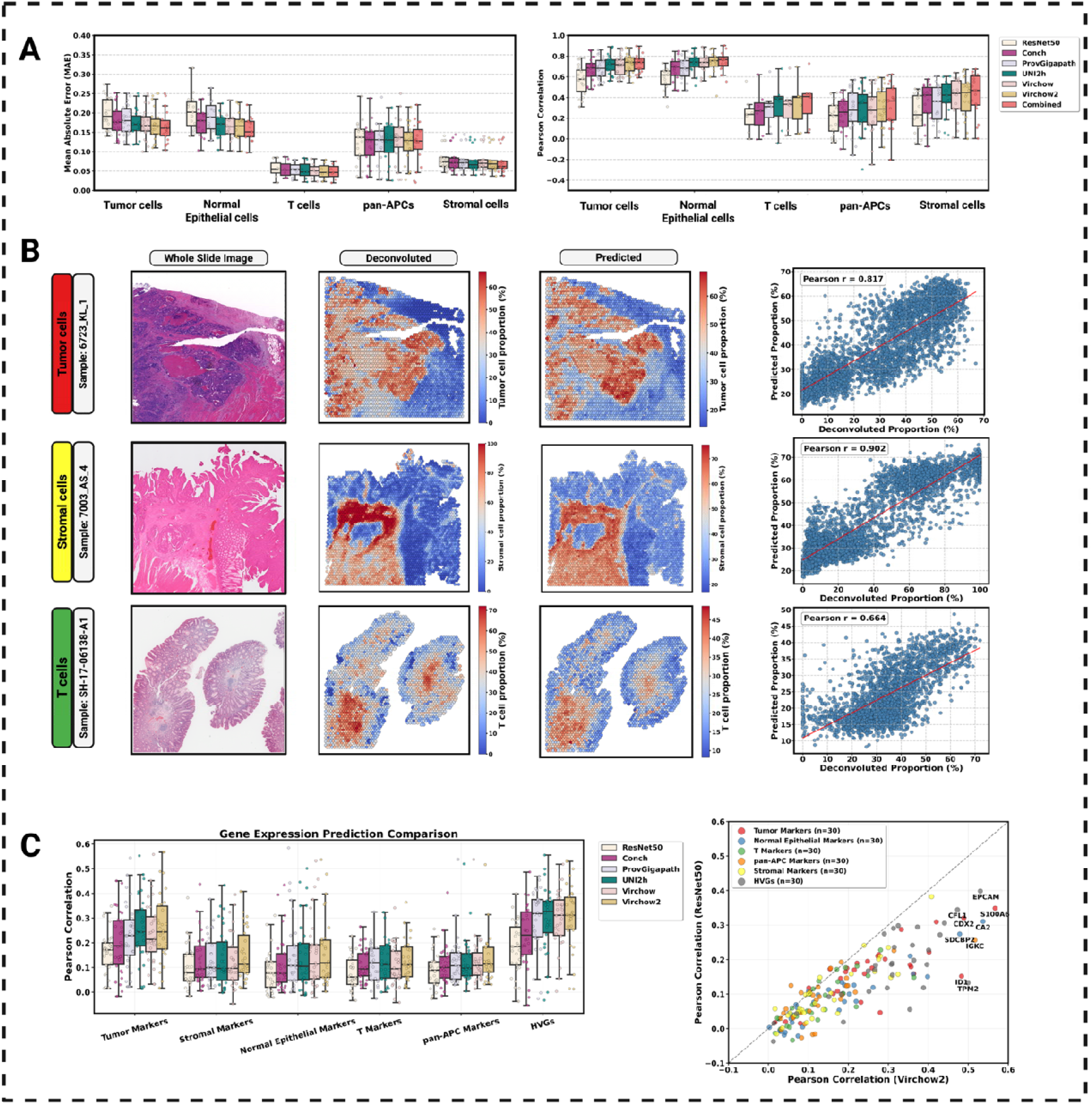
Cross-validated performance, gene expression, and spatial agreement of STpath predictions for colorectal cancer. (A) Leave-one-individual-out (LOIO) performance across 5 patholog foundation AI models, reference model and the combined model for 5 major cell-type categories. Left: Mean Absolute Error (MAE). Right: Pearson correlation. Each point represents an individual. Boxe show interquartile ranges with medians. (B) Representative whole-slide results for Tumor, Stromal, and T-cell enrichment. Columns show the original H&E WSI, deconvoluted proportions from spatial transcriptomics, LOIO model-predicted proportions, and spot-wise scatter plots comparing prediction with deconvoluted values. (C) Gene expression prediction comparison. Left: Pearson correlations between predicted and true gene expression across 5 marker gene sets (30 genes each) and 30 HVGs. For each gene, Pearson correlation was computed under the LOIO strategy and the median across individuals was taken. Each point represents one gene. Right: Scatter plot comparing ResNet50 and Virchow2, with each dot representing a single gene.

Because MAE is influenced by the average abundance of each cell type (Figure 2C), we primarily relied on Pearson correlations for comparisons across cell types. Prediction performance was highest for tumor and stromal cells, with the combined model achieving correlations exceeding 0.7 for both. In contrast, correlations were lower for T cells and normal epithelial cells (median LOIO Pearson correlations ∼0.4), likely reflecting their low proportions in many samples (Figure 2C). The robust prediction performance of STpath on unseen patients is illustrated on three representative H&E WSIs: tumor-enriched, stromal-enriched, and T-cell-enriched samples (Figure 4B). Although predictions showed a trend of regression towards the mean, they nonetheless captured tissue domains dominated by a single cell type.

### STpath Accurately Estimates Gene Expression for a Subset of Genes

We next evaluated the ability of STpath to predict gene expression (Figure 4C). For this analysis, we curated a panel of 180 genes, including 150 cell type–specific marker genes (30 per cell type) and 30 highly variable non-marker genes (HVGs). Across all models, ResNet50 demonstrated the weakest predictive performance, with median Pearson correlations of ∼0.2 for HVGs and even lower median Pearson correlations for cell type–specific marker genes. This trend aligns with recent observations from GHIST^19^, which also used a ResNet-based architecture. By contrast, all pathology foundation models outperformed ResNet50, with Virchow2 achieving the best overall accuracy (Figure 4C).

Performance varied markedly across individual genes. Some genes, such as the colorectal cancer marker **S100A6**, were highly predictable, reaching a LOIO Pearson correlation of 0.57 under Virchow2 (Figure 4C, right; Figure S1C), whereas some other colorectal cancer markers showed correlations below 0.1. These findings highlight that predicting cell type proportions is inherently more reliable than predicting expression of individual genes (Figure 2D). Consistently, comparing Figures 4A and 4C shows that Pearson correlations for cell type proportion prediction exceed those for gene expression prediction.

We also computed partial correlations between predicted and observed gene expression while conditioning on cell type proportions. These partial correlations were uniformly lower than the unadjusted correlations (Figure S1D), indicating that a substantial fraction of the predictive signal arises from underlying cell type compositions.

### Multi-resolution Robustness and Quantitative Spatial Segmentation

STpath enables systematic analysis of cell type composition across multiple spatial resolutions. By default, cell type proportions are estimated for each ∼256 × 256-pixel tile, resulting in an approximately 70 × 70 grid for a typical H&E whole-slide image. To evaluate robustness to spatial scale, we randomly selected representative slides and computed average cell type proportions while varying the grid resolution from 40 × 40 to 100 × 100. Across all resolutions, estimated cell type compositions remained highly stable, with fluctuations not exceeding 8% (Figure 5A). This robustness was further demonstrated in representative samples enriched for tumor cells, stromal cells, and T cells, respectively (Figure 5B, Figure S4). Across all cases, global spatial patterns of cell type composition were preserved across resolutions, while finer grids provided increased spatial detail.

**Figure 5.**
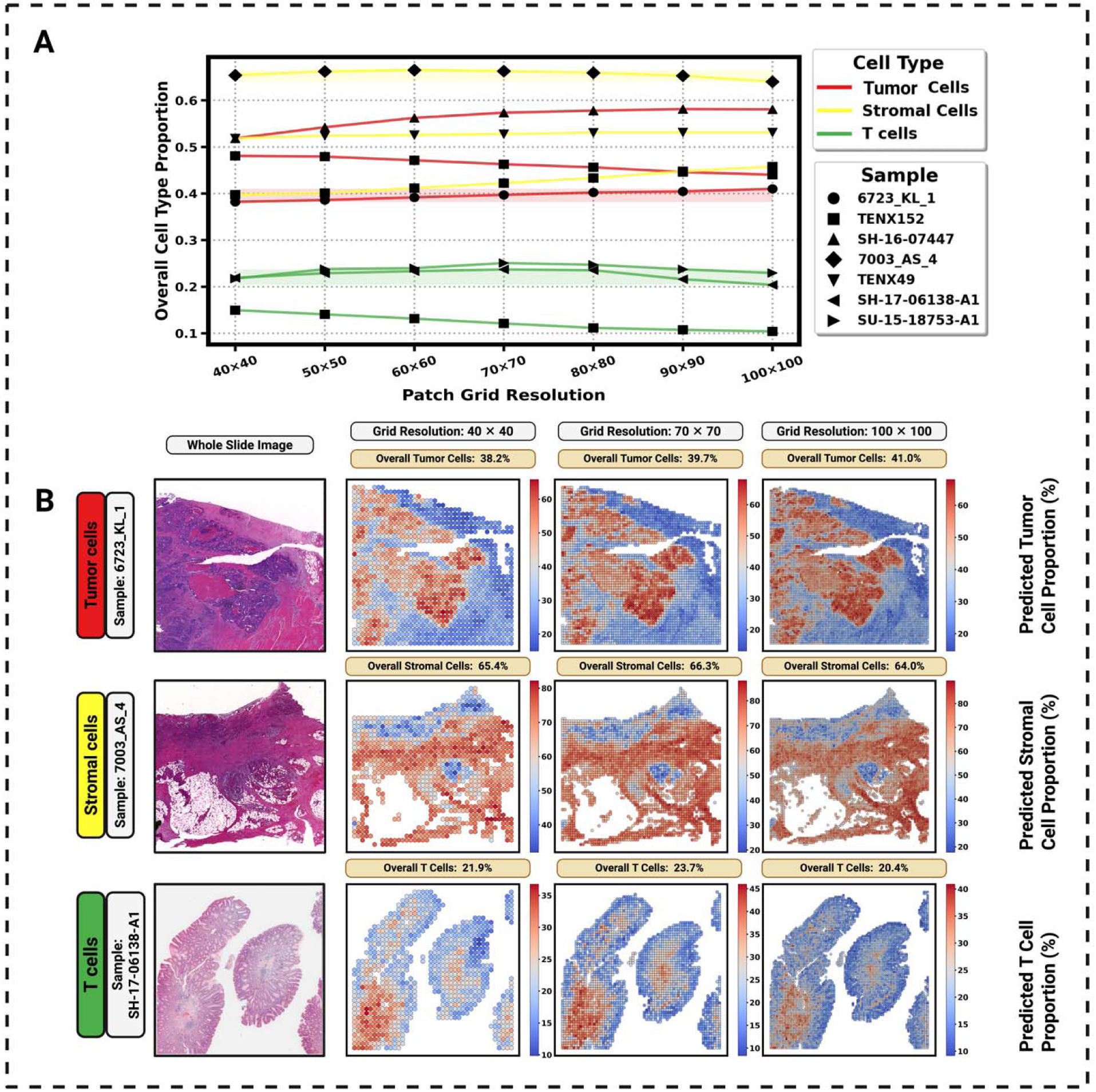
Robustness of overall cell type proportion estimates to patch-grid resolution. (A) Overall proportions for Tumor cells, Stromal cells, and T cells remain stable as the patch grid changes from 40×40 to 100×100 across multiple representative samples. (B) Three example WSIs (Tumor-, Stromal-, and T-cell–enriched cases) visualized at 40×40, 70×70, and 100×100 grids. Hexagonal binned heatmaps show patch-level predicted proportions, with slide-level overall proportions annotated above each map.

Beyond multi-resolution stability, the quantitative outputs of STpath enable spatial segmentation at different cell type proportion cutoffs. Increasing the threshold progressively restricted selected regions, enabling precise localization of highly enriched areas. For example, applying a >50% threshold for tumor cell proportion yielded spatial maps that clearly delineated regions with substantial tumor enrichment (Figure S6).

Together, these results demonstrate that STpath provides a robust and flexible framework for tile-level cell type quantification, enabling consistent analysis across spatial scales and quantitative definition of enriched tissue regions.

### Using STpath to Identify Prognostic Tumor Microenvironment Signatures for Colorectal Cancer

We demonstrated the generalizability and clinical utility of trained STpath XGBoost models on an external dataset of H&E images from 232 in The Cancer Genome Atlas Colon Adenocarcinoma (TCGA-COAD) Collection. STpath gave the predicted cell type proportions of each image tile (Figure 6A, top row). To facilitate explanation and down-stream analysis, we labeled each tile by its composing cell types. Each tile was assigned a label of one cell type if its predicted proportion exceeded a prespecified threshold, and a tile could be labeled by multiple cell types (Figure 6A, bottom row). See Method Section for details on the threshold choices for each cell type.

**Figure 6.**
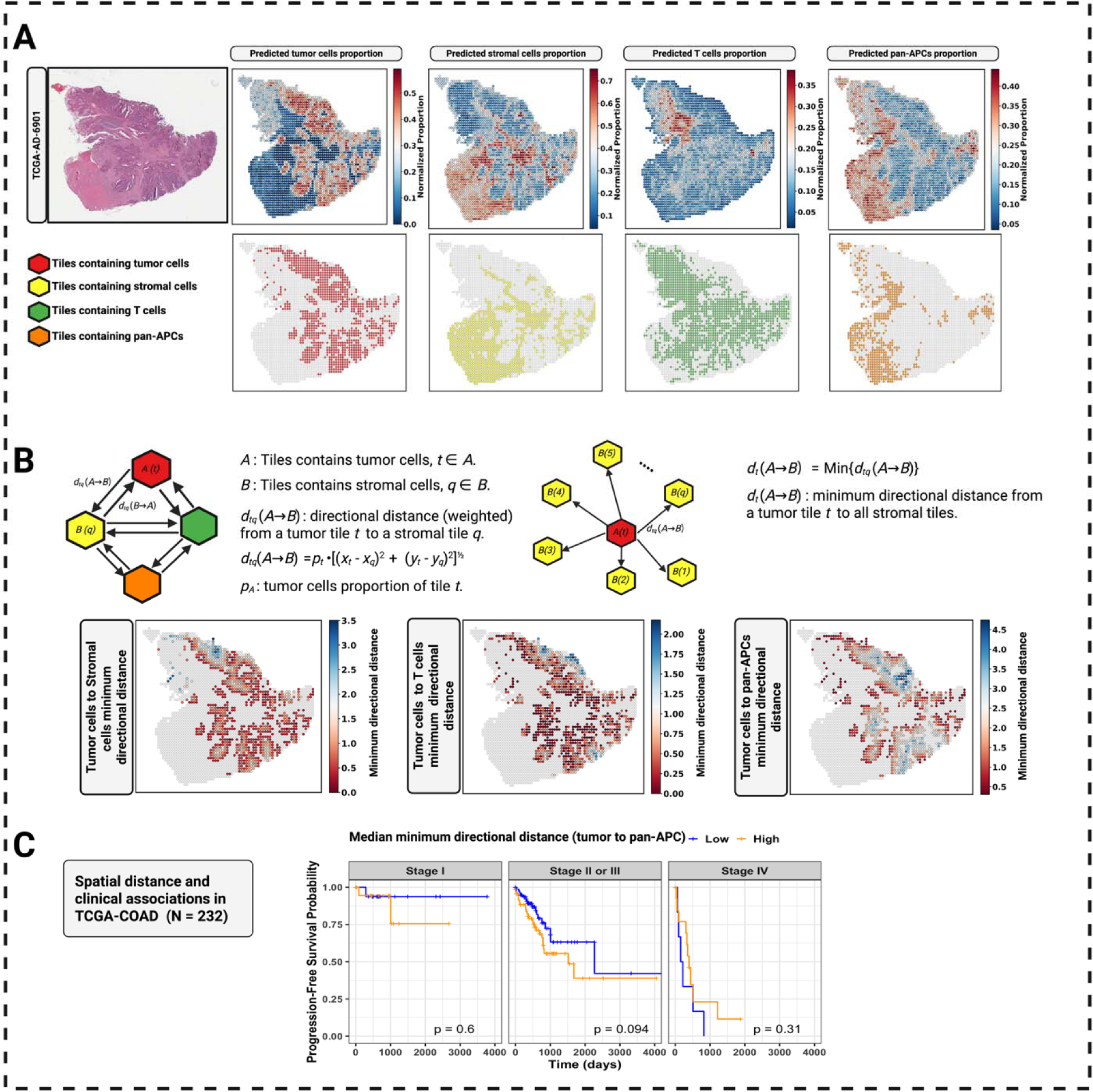
STpath predictions enable spatial distance metrics and clinical associations in TCGA-COAD. (A) Cell type prediction and tile classification. Top row: Predicted cell type proportion (normalized) for tumor cell, stromal cell, T-cells, and pan-APCs in a representative TCGA-COAD sample. Bottom row: Tile classification based on proportion thresholds (tumor cell>30%, stromal cell >40%, T-cells >10%, pan-APCs >30%). (B). Minimum directional distance calculation. Schematic illustrating calculation of weighted minimum distance from a tumor tile to stromal tiles, where distance is weighted by the source tile’s cell proportion. Bottom row: Spatial distribution of minimum directional distances from tumor-to-stromal cell, T-cells, and pan-APCs. (C) Kaplan-Meier curves for progression-free interval (PFI). Left: PFI stratified by median minimum directional distance groups (low vs. high) for tumor-to-pan-APCs, stratified by AJCC tumor stages. N=232 TCGA-COAD patients. P-values from log-rank test.

Given cell type labels for each image tile, we could calculate the distances between tiles harboring any two cell types and we sought to use the summary of such distance metrics to derive spatially-resolved prognostic features. For a tile *t* having a label of cell type *A*, we calculated a directional distance from source cell type *A* to target cell type *B*, denoted by *d_t_* (*A* → *B*) as the minimum Euclidean distance from tile *t* to any tile having the label of cell type *B*. (Figure 6B). We adopt the minimum distance to measure whether cell type *A* in this tile can interact with any cells of type *B*. Of course, if this tile has both labels of *A* and *B*, this distance is zero. An illustration of this directional distance was shown in Figure 6B, where the tumor cells in a small region on the top of the slide were further away from stromal cells.

Next, we summarize the directional distances *d_t_* (*A* → *B*) across all tiles by its median, which measures the typical distance from cell type *A* to cell type *B* across all tiles having cell type *A*. The median is highly concordant with other summary metrics (mean, standard deviation, and interquartile range) (Figure S5), therefore we use the median as a robust representative statistic in downstream analyses. Focusing on the directional distances from tumor cells to immune cells in TCGA-COAD samples, we found shorter tumor-to-pan-APC distances were associated with better survival outcomes, either before (p-value 0.029, HR 1.15, 95% CI: 1.014-1.304) or after adjusting for age, sex and tumor stage (p-value 0.027, HR 1.16, 95% CI: 1.017-1.321). Shorter tumor-to-T-cell distances and tumor-to-stromal distances were also associated with better survival time after adjusting by age, sex, and tumor stage (Figure 6C, Table 1).

**Table 1.** Cox proportional hazards regression models of median distance features for progression-free survival in TCGA-COAD. Hazard ratios with 95% confidence intervals and *p*-values are reported from the Cox proportional hazards model (either unadjusted or adjusted) with progression-free survival as the outcome. Significant associations (p < 0.05) are shown in bold.

| Median Distance | Unadjusted Cox Model |  | Adjusted Cox Model |  |
| --- | --- | --- | --- | --- |
| | HR (95% CI) | $p$-value | HR (95% CI) | $p$-value |
| Tumor cells to Stromal cells | 1.137 (0.944–1.368) | 0.176 | 1.226 (1.010–1.489) | **0.039** |
| Tumor cells to Pan-APCs | 1.150 (1.014–1.304) | **0.029** | 1.159 (1.017–1.321) | **0.027** |
| Tumor cells to T cells | 1.021 (0.995–1.048) | 0.112 | 1.029 (1.000–1.058) | **0.047** |
| Stromal cells to Tumor cells | 0.940 (0.822–1.075) | 0.363 | 0.938 (0.825–1.067) | 0.329 |
| Stromal cells to Pan-APCs | 1.095 (0.981–1.224) | 0.107 | 1.068 (0.936–1.218) | 0.328 |
| Stromal cells to T cells | 1.008 (0.983–1.033) | 0.544 | 1.010 (0.983–1.038) | 0.474 |
| Pan-APCs to Tumor cells | 1.025 (0.908–1.156) | 0.692 | 1.054 (0.935–1.188) | 0.387 |
| Pan-APCs to Stromal cells | 0.935 (0.707–1.236) | 0.637 | 0.935 (0.688–1.272) | 0.669 |
| Pan-APCs to T cells | 1.011 (0.974–1.050) | 0.566 | 1.016 (0.975–1.060) | 0.445 |
| T cells to Tumor cells | 0.954 (0.685–1.329) | 0.781 | 1.067 (0.759–1.500) | 0.709 |
| T cells to Stromal cells | 0.987 (0.684–1.424) | 0.943 | 1.095 (0.748–1.603) | 0.642 |
| T cells to Pan-APCs | 0.978 (0.781–1.224) | 0.846 | 1.047 (0.831–1.321) | 0.695 |

Tumor mutation burden is an important molecular characteristic of colorectal cancer. A subset of colorectal cancer has a high number of somatic point mutations and is often referred to as hyper-mutated subtype of colorectal cancer. Patients with hypermutated cancer are more likely to respond to cancer immunotherapy^23^, making the relationship between tumor mutation burden and tumor microenvironment of particular interest. Understanding this association may help elucidate mechanisms underlying immunotherapy response. Our results showed that shorter tumor-to-pan-APC (antigen presenting cells) and tumor-to-T-cell distances are associated with higher mutation burden (Table 2). These findings are consistent with previous studies^24^ and further supports the biological relevance and utility of STpath models.

**Table 2.**
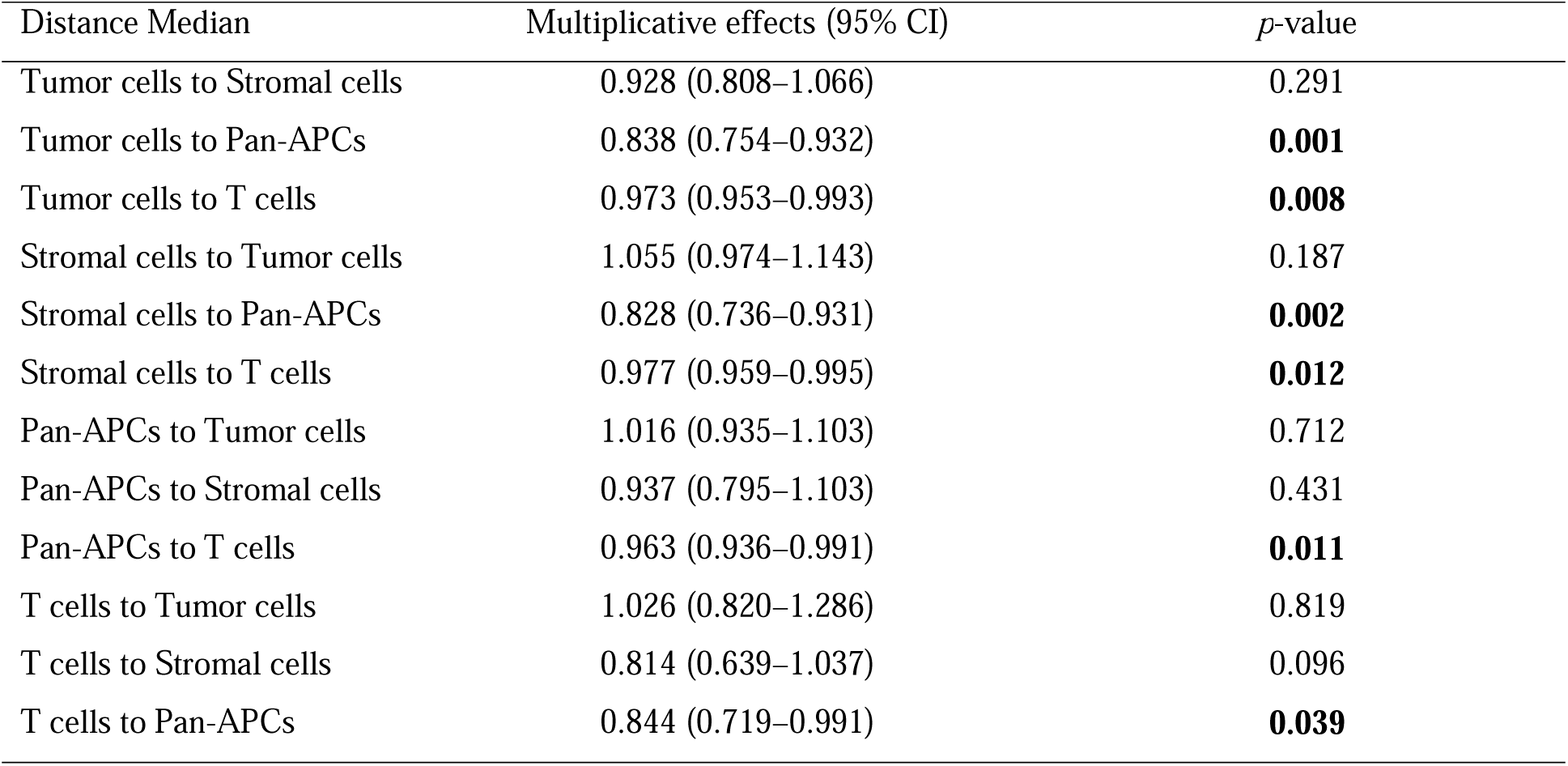
Adjusted log-linear regression model of median distance features for mutation burden in TCGA-COAD. Multiplicative effects with 95% confidence intervals and *p*-values are reported from the joint model with log-transformed somatic mutation burden as the outcome. Predictors include AJCC tumor stage status, age, gender, and each of the spatial distance medians. Significant associations (p < 0.05) are shown in bold.

### STpath Models Should Be Trained Separately for Each Cancer Type

To evaluate the generalizability of STpath in other cancer types and assess whether different cancer types require different fine-tuned foundation models (i.e., foundation model + XGBoost), we trained STpath models for breast cancer. We defined five major cell types for breast cancer, consistent with those used for colorectal cancer, and estimated cell type composition for each image tile using paired spatial transcriptomic data (Figure S7A-B). The estimated cell type proportions are strongly correlated with relative marker gene expression, confirming the accuracy of the cell type proportion estimates (Figure S7C-D).

We trained XGBoost models to predict cell type proportions using embeddings from each of the five pathology foundation models as well as from ResNet50. Similar to our observations in colorectal cancer, prediction performance was highest for tumor epithelial cell proportions (Figure 7A, Figure 4A). However, stromal cell prediction in breast cancer was notably less accurate than that observed in colorectal cancer (Figure 7A, Figure 4A). Interestingly, breast cancer samples exhibited relatively strong predictive performance for pan-APC populations, with a median Pearson correlation exceeding 0.6 (Figure 7A). The combined model integrating features from all five pathology foundation models consistently achieved the best overall performance, while Virchow2 performed comparably well across multiple cell types (Figure 7A). The normal epithelial cells remained the most challenging cell type to predict, consistent with findings in colorectal cancer (Figure 7A, Figure 4A). Across all cell types, pathology foundation models substantially outperformed the ResNet50 baseline (Figure 7A), highlighting the advantage of histopathology-specific pretrained representations over generic computer-vision backbones.

**Figure 7.**
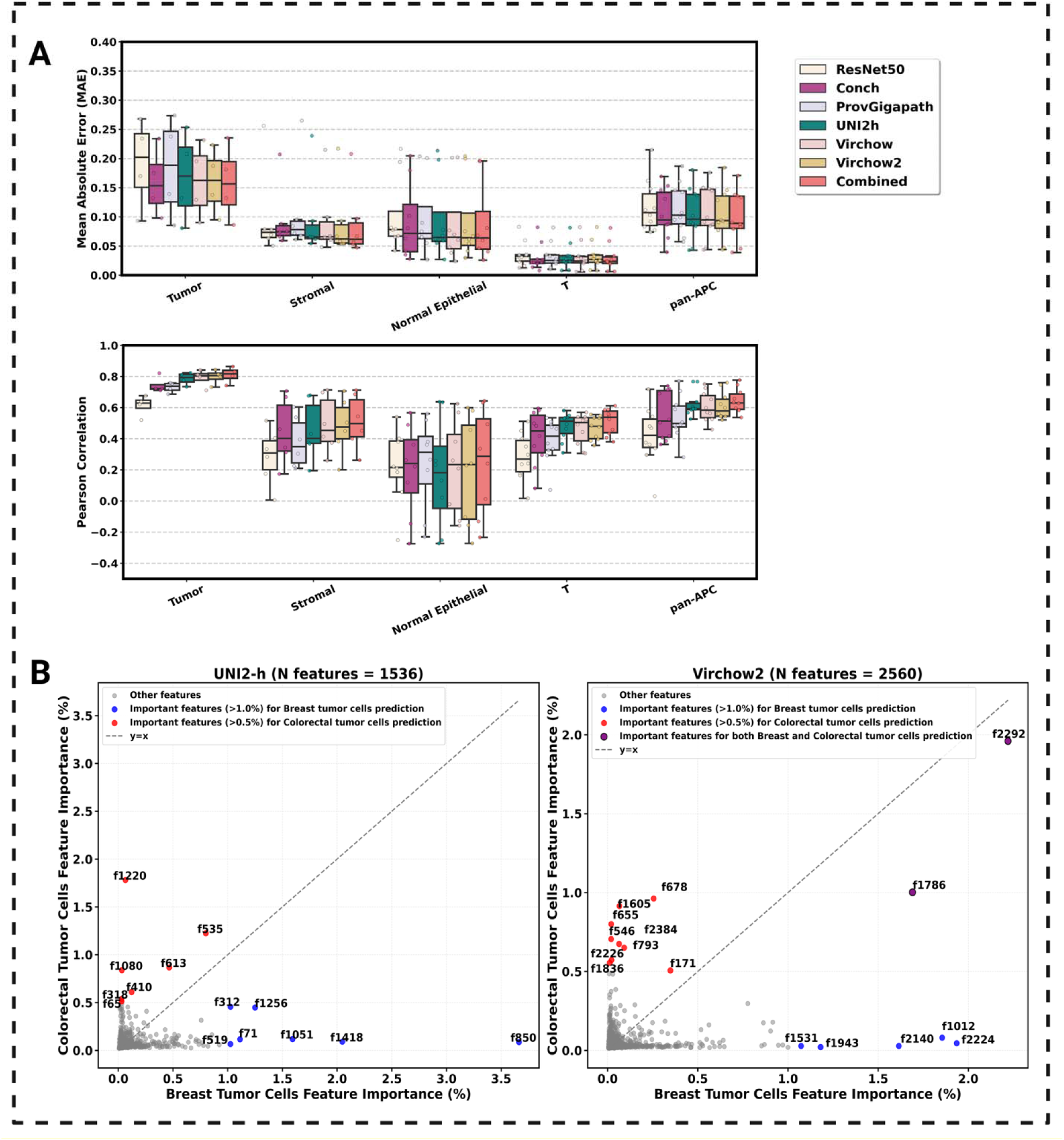
Cross-validated performance for breast cancer, feature importance comparison between breast colorectal cell and breast tumor cell prediction. (A). LOIO cross-validation performance. Model performance across five pathology foundation AI models, a reference model, and a combined model for eight major cell-type categories. Left: MAE. Right: Pearson correlation. Each point represents an individual sample. Boxes show interquartile ranges with median lines. (B) Feature importance comparison between colorectal and breast tumor cell prediction models. Scatter plot comparing XGBoost feature importance for tumor cell prediction between BRCA and COAD. Left: UNI2h (N features = 1536). Right: Virchow2 (N features = 2560).

A key practical question is whether distinct STpath models are required for different cancer types. To address this, we compared feature importance for tumor cell prediction between colorectal and breast cancer (Figure 7B). It was clear that feature importance differed markedly between the two cancers. For example, under UNI2-h, feature #850 was among the most informative predictors for breast cancer tumor cells but contributed minimally to colorectal cancer, whereas feature #1220 showed the opposite pattern, highly informative in colorectal cancer but unimportant in breast cancer (Figure 7B). Similar patterns were also observed for Virchow2; however, we also identified a small number of shared features, such as feature #2292 of Virchow2, which exhibited high importance in both breast and colorectal cancer tumor cell proportion prediction models (Figure 7B). Therefore, we concluded that STpath XGBoost models need to be trained for different cancer types separately.

## Discussion

In this study, we present STpath, a cancer-specific framework that enables the translation of abstract histopathology foundation model embeddings into biologically interpretable and clinically actionable features. STpath does not retrain foundation models; instead, it learns cancer-specific mappings from pretrained embeddings to molecular or cellular features, enabling interpretability and reducing technical artifacts.

A notable finding of our work is that all evaluated foundation models exhibit pronounced batch effects at the image level. The tiles from the same image frequently cluster together in embedding space, even when they differ markedly in cellular composition. This phenomenon likely reflects image-specific characteristics such as staining intensity, tissue processing, and/or scanner-specific artifacts. This batch effect confounds tile-level biological interpretation. Interestingly, supervised feature selection via XGBoost can substantially mitigate these batch effects by extracting information most relevant to specific targets, e.g., cell type composition or gene expression.

We find that the leave-one-individual-out (LOIO) evaluation strategy is essential for obtaining an objective assessment of prediction accuracy. Because tiles extracted from the same image may share substantial technical and biological similarity, randomly splitting the tiles into training and testing sets allows tiles from the same image to appear in both sets, leading to information leakage. Such leakage can substantially inflate apparent performance and does not reflect true generalization to unseen patients. Consistent with this concern, when we applied a random tile-level split as an alternative evaluation strategy, we observed markedly higher, but misleading, prediction accuracies (Figure S8).

Our systematic comparison of five state-of-the-art pathology foundation models reveals that these foundation models trained on large, diverse histopathology data consistently outperform a generic ResNet50 backbone for predicting cell type composition and gene expression. However, no single foundation model dominates across all cell types. Instead, different models preferentially capture distinct morphological information, resulting in complementary predictive strengths. An STpath model that integrates features across all foundation models achieves improved accuracy, highlighting the value of ensemble representations for complex tissue microenvironments.

Although pathology foundation models are designed to be broadly applicable, our analyses show that the mapping from image embeddings to biological features is cancer-type specific. Feature importance profiles differed markedly between colorectal and breast cancer, with features informative in one cancer often contributing little in another. These results indicate limited transferability across cancer types and justify the need for cancer-specific STpath models to achieve accurate and interpretable biological inference.

STpath enables consistent annotation of H&E-stained images in large studies, and such annotations can be used in downstream association studies, as illustrated in our TCGA colon cancer study presented herein. STpath could also be an assistant to a pathologist, though it cannot replace a pathologist’s expertise in clinical settings. In a few testing examples where we have obtained pathologist’s manual annotations, STpath’s annotation for tumor regions are highly consistent with manual annotation (AUC > 0.9), but its annotation for stromal regions is less accurate (AUC = 0.86), likely because stromal is a broad cell type and it includes many subcategories (Figure S9). Future studies with larger amount of training data could further improve STpath’s ability to dissect different subtypes of stromal cells.

Several limitations merit consideration. First, STpath relies on spatial transcriptomics data for supervised model training, which may limit applicability in tumor types with sparse datasets. At the time of preparing this paper, most available spatial omics data were spot-level, such as 10X Visium. Studies combining H&E images and cellular-resolution spatial omics (e.g., 10X Xenium) are emerging, though they remain limited in sample size^19^ and primarily focus on within-sample prediction^20^. We expect that future work to expand STpath model training on large scale Xenium data will further improve its performance. Second, gene expression prediction remains challenging, particularly for genes with low expression variation. Finally, although we demonstrate spatially resolved prognostic features can be found, much more work is needed to identify more informative biomarkers for cancer prognosis or possibly treatment outcome prediction.

In summary, STpath establishes a general strategy for transforming pathology foundation model embeddings into interpretable, spatially resolved features tailored to specific cancer types. By addressing batch effects, leveraging complementary foundation models, and grounding predictions in spatial transcriptomics, STpath advances the safe and effective translational use of AI in digital pathology. As foundation models continue to evolve, frameworks such as STpath will be essential to ensure that their power is harnessed in biologically meaningful and clinically responsible ways.

## Methods

### Data Preparation

For colorectal cancer, the training cohort included 63 H&E images (with paired Visium data) from three datasets: 32 images from the Colon Molecular Atlas Project (Vanderbilt HTAN^25–27^), 13 images from HEST-1K^28^ dataset, and 18 images from an in-house Fred Hutch dataset (Yu et al., unpublished data). It is important to note that the samples in HEST-1K originally included part of the Colon Molecular Atlas Project; in our study, we used the HEST-1K colorectal subset excluding the Colon Molecular Atlas Project to avoid redundancy. For applications using external H&E image data, we used H&E images and corresponding clinical information from the TCGA-COAD collection^29–30^. For breast cancer, the training cohort includes 11 H&E images (with paired Visium data) assembled from multiple publicly available sources, including five images from the 10x Genomics website, four images from Liu et al.^31^, and two images from Wu et al.^32^.

H&E WSIs were processed to create tissue tiles aligned with spatial transcriptomics coordinates. For each sample, we first established coordinate alignment between WSIs and spatial transcriptomics data by calculating scaling factors based on the full-resolution pixel coordinates and WSI dimensions. Sample-specific geometric transformations (rotation, mirroring) were applied as needed to ensure proper spatial registration.

We implemented Otsu automatic thresholding^33^ for tissue region identification. Each WSI was converted to grayscale and segmented using Otsu’s method, where tissue regions (darker areas in H&E staining) were identified by applying the threshold:

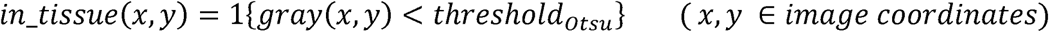

Here, *gray*(*x*, *y*) denotes the grayscale intensity at pixel (*x*, *y*), with values from 0 to 255 and larger values corresponding to white background. *in_tissue* (*x*, *y*) is the binary segmentation radius 15, removal of small regions (< 2,000 pixels), followed by dilation (disk radius 10) and mask. Morphological operations were performed to refine segmentation results: closing with disk erosion (disk radius 8) to connect nearby tissue regions while preserving overall morphology.

Connected component labeling was applied to identify distinct tissue regions. Spatial transcriptomics spots were assigned to tissue regions based on their scaled coordinates. Image tiles were extracted at spatial transcriptomics coordinates using median inter-spot distances to determine optimal patch radius. Tiles with excessive white space (>40%) and regions with < 30 valid tiles were excluded to ensure tissue coverage and statistical reliability. Detailed patch extraction procedures are provided in Supplementary Methods: Spatial Transcriptomics Data Integration.

We used scRNA-seq data from Heiser et al.^26^ and from Mo et al.^34^ as reference for cell type deconvolution for colorectal cancer and breast cancer, respectively. Marker genes were identified through differential expression analysis using Wilcoxon rank-sum tests with multiple comparison correction. A total of 899 and 577 marker genes were identified across cell types for colorectal cancer and breast cancer, respectively (Figure S9 and Figure S10). See Supplementary Methods: Single-cell RNA-seq Reference Data Processing for detailed procedures.

### Cell Type Deconvolution and Results Validation

Cell type deconvolution for each spot in Visium data was performed by CARD with a minor modification to the reference expression matrix. By default, CARD applies its own informative gene selection procedure when a genome-wide scRNA-seq reference matrix is provided. In our analysis, we disabled this step and instead directly supplied the gene expression data matrix of the pre-selected cell type marker genes. This modification led to a slight improvement of deconvolution accuracy as evaluated in the next step.

To validate the accuracy of our cell type deconvolution results, we implemented a validation approach that compared predicted cell type proportions with the expression of cell type-specific marker genes, which were identified by differential expression analysis of the scRNA-seq reference dataset. Gene expression counts were normalized by library size, followed by CARD signature matrix normalization to account for baseline expression differences across cell types. For each cell type, we computed the relative marker gene expression (RMGE) as a proxy for cell type abundance, providing a standardized metric for validating CARD deconvolution results. See Supplementary Methods: Deconvoluted Results Validation for detailed procedures.

For colorectal cancer, we considered five major cell type categories: Tumor cells, Normal epithelial cells, T cells, pan-antigen-presenting cells (pan-APCs), and Stromal cells. Here pan-APCs include B cells, myeloid cells and plasma cells. The proportion of these five cell types was subsequently used for constructing the predictive models (see Figure S11 for details). For breast cancer, we similarly considered the same five cell type categories; however, the tumor cell group consisted of distinct breast cancer subtype groups.

### Foundation Model Features Extraction and Predictive Models Building

We employed five pathology foundation models to extract numerical embeddings from image tiles (Figure S3): Conch^3^, ProvGigapath^4^, UNI2h^5^, Virchow^6^, and Virchow2^7^. We also included ResNet50 pretrained on ImageNet as a baseline vision model. For each foundation model, H&E image tiles (224×224 pixels) underwent model-specific preprocessing including resizing, center cropping, and ImageNet normalization. Embeddings were extracted using batch processing (batch size = 32) with no gradient computation to ensure computational efficiency. The dimension of the extracted embeddings (i.e., the number of features, denoted by *p*) varied across models: Conch (*p* = 512), Prov-GigaPath (*p* = 1536), UNI2-h (*p* = 1536), Virchow (*p* = 2560), and Virchow2 (*p* = 2560). For Virchow and Virchow2 models, we concatenated the class token with the averaged patch tokens to obtain the final feature representation.

For each cell type and foundation model combination, we trained XGBoost regression models to predict cell type proportions. Model hyperparameters were optimized through grid search and configured as follows: learning rate of 0.01, maximum tree depth of 12, minimum child weight of 5, column sampling ratio of 0.05, subsample ratio of 0.5, *L*_1_ regularization of 0.1, *L_2_* regularization of 0.01, and histogram-based tree construction method for computational efficiency. The objective function was set to be squared error for regression tasks.

We implemented a rigorous LOIO cross-validation strategy to evaluate model generalizability. For each individual in the dataset, we trained models using the image tiles from all other individuals and tested on this held-out individual. This approach ensures that the model evaluation reflects real-world performance on unseen patients, avoiding data leakage that could occur with random train-test splits. We trained separate XGBoost regression models for each combination of cell type and foundation model, resulting in 30 models (5 cell types × 6 models including ResNet50 baseline) for both colorectal and breast cancer. We calculated feature importance scores across all folds using XGBoost built-in feature importance metric (Gain). For each cell type, we averaged importance scores across all cross-validation folds and ranked features by their mean importance. Important features were selected using a cumulative importance threshold of 30%, ensuring we retained the most predictive features while reducing dimensionality. This resulted in model-specific feature sets (excluding ResNet50): Conch (154), Prov-GigaPath (461), UNI2-h (461), Virchow (768), and Virchow2 (768). Details are in the Supplementary Files 3-4. To leverage complementary information from different foundation models, we created combined models by concatenating the selected important features from all five foundation models, yielding five combined models for the five cell types of both colorectal cancer and breast cancer.

Model performance was evaluated using two metrics: Mean Absolute Error (MAE) to assess prediction accuracy, and Pearson correlation coefficient to measure the linear relationship between predicted and true cell type proportions.

To quantify the complementary contributions of different foundation models, we conducted a dual-metric analysis to evaluate both absolute and normalized feature contributions. For each cell type, XGBoost models were trained using concatenated features from all five foundation models (UNI2-h, Virchow, Virchow2, Prov-GigaPath, and Conch), with feature importance scores computed through cross-validation. Absolute contribution measured the raw predictive power of each model, while normalized contribution accounted for varying feature dimensions across models, revealing per-feature efficiency.

Complementarity analysis employed pairwise Ridge regression to assess feature dependence between foundation models, with lower *R*^2^ values indicating higher complementarity. This approach demonstrated that different foundation models captured distinct morphological aspects, with some providing broad tissue architecture information while others contributing highly discriminative features for specific cell types. See Supplementary Methods: The Complementary Analysis of Foundation Models for detailed procedures.

Computational Performance. All experiments were conducted on an Apple MacBook Pro equipped with an M3 Max chip (14-core CPU, 30-core GPU, 64GB unified memory) using PyTorch with Metal Performance Shaders (MPS) acceleration. The framework also supports NVIDIA CUDA GPUs and CPU-only execution. Feature extraction for approximately 330,000 image tiles (256×256 pixels) using batch size 32 required approximately 4-6 hours per foundation model. Peak memory usage ranged from 4-12 GB depending on feature dimensionality, with combined models requiring up to 16 GB. The complete pipeline, including feature extraction and training for all cell type-model combinations, can be completed within 240 hours on a single GPU-enabled workstation.

### Gene Expression Prediction Analysis

We implemented a gene expression prediction pipeline using all five foundation models’ and ResNet50’s features from a merged and deduplicated dataset combining all 5 cell-type-specific training sets for colorectal cancer. A curated set of 180 predictor genes was selected, comprising: (1) 150 cell type-specific marker genes (30 per cell type: Tumor cells, Normal epithelial cells, T cells, Stromal cells, pan-APCs) identified through differential expression analysis with duplicate genes across cell types removed to ensure specificity; (2) an additional 30 highly variable genes selected exclusively from non-marker genes. Using the same LOIO cross-validation, we trained separate XGBoost regression models for each gene with the same optimized hyperparameters.

To quantify the extent to which gene expression predictions are mediated by cell type composition, we employed partial correlation analysis controlling for cell type proportions. Specifically, for each individual, we calculated both regular Pearson correlation between predicted and actual gene expression and partial correlations that control the confounding effects of cell type composition.

Cell type proportions were transformed using centered log-ratio (CLR) transformation^35^ to address the compositional nature of the data. This transformation converts the compositions to real valued vectors ranging from -inf to inf:

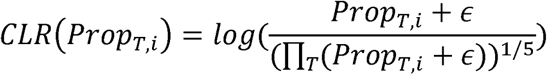

where *Prup_r,i_* is the cell type proportion of cell type *T* in tile *i*, and ∈ = 1 x 10^-6^ prevents numerical instability.

For each gene, the partial correlation was computed by regressing both predicted and actual gene expression against CLR-transformed proportions and calculating correlations between the resulting residuals:

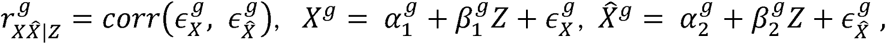

where *x^g^* denotes the observed (log-normalized) gene expression for gene g, *̂X^g^* denotes the XGBoost-predicted gene expression, £ denotes the CLR-transformed cell type proportions, 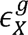 and 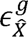 are the residuals after adjusting for £.

### Cell Type Proportion Consistency Assessment Across Different Image Tile Sizes

For each H&E WSI, we assumed equal cell density across valid tissue regions and calculated overall proportions as the arithmetic mean of individual tile predictions:

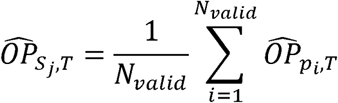

where 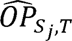 denotes the estimated overall proportion of cell type *T* for the sample *j*, 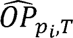 is the predicted proportion of cell type *T* in tile *i*, and *N_valid_* is the number of valid tissue tiles.

To assess model stability and robustness to different image processing conditions, we implemented a comprehensive consistency analysis using varying patch segmentation strategies. For three representative samples across different cell types (cancer cells, stromal cells, and T cells), we tested seven different grid resolutions (40×40, 50×50, 60×60, 70×70, 80×80, 90×90, and 100×100 tiles per WSI). Each segmentation was processed through the complete STpath pipeline: foundation model feature extraction, cell type proportion prediction using trained LOIO XGBoost models, and overall cell type proportion calculation. Consistency was quantified by comparing overall cell type proportion estimates across different grid sizes, with stable predictions indicating robust model performance.

### Quantitative Tissue Segmentation

We implemented a probabilistic soft segmentation approach to visualize tissue heterogeneity based on predicted cell type enrichment. Each whole slide image was divided into a 100×100 grid of tiles, and the tiles exceeding predefined cell type-specific proportion thresholds were highlighted with transparent color overlays. Threshold values were empirically determined based on biological relevance. This approach enables pathologists to identify regions of interest and assess spatial tissue composition patterns across different scales.

### Statistical Modeling for Clinical Associations-TCGA Colorectal Cancer Cohort Analysis

We applied STpath to 232 TCGA-COAD samples and 431 The Cancer Genome Atlas Breast Invasive Carcinoma (TCGA-BRCA) samples to evaluate its clinical utility. Although the TCGA repository contains a larger number of H&E slides, we performed manual quality filtering to ensure reliable spatial analyses. Slides with excessively small tissue regions, visible staining or scanning artifacts, contamination, or lens-related blurring were excluded prior to downstream processing. Valid tissue tiles were classified into cell-type categories using study-specific proportion thresholds. For each valid tile, features were extracted using pretrained pathology foundation models, and cell type proportions were predicted using pre-trained XGBoost models. To calibrate predictions, raw predicted proportions were mapped to the validation set distribution using quantile mapping, where the validation proportions were scaled to [0, 1] by setting values above the 95th percentile to 1. The calibrated proportions for all five cell types were then renormalized to sum to 1 for each tile. For TCGA-COAD, the thresholds are tumor (>30%), stromal (>40%), T cells (>10%), and pan-APCs (>30%). For TCGA-BRCA, the thresholds were: cancer epithelial (>15%), pan-APCs (>30%), stromal (>30%), and T cells (>30%). These thresholds were determined empirically based on the distribution of calibrated normalized proportions across all tiles, corresponding approximately to the 25th percentile for each cell type, thereby selecting tiles with above-average abundance of the corresponding cell population. Because tiles could contain mixtures of cell types, a tile could be assigned to multiple cell types when the predicted proportions exceeded the corresponding thresholds. Tiles that met none of the thresholds were excluded from spatial analyses. For spatial analysis, minimum directional Euclidean distances were computed between cell type pairs. For two classes *A* and *B*, and each tile t of class *A*, and any tile *q* of class *B*, the distance weighted by cell type proportion from tile t to *q* was defined as:

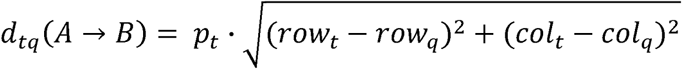

where *p_t_* is the class *A*’s cell type proportion of tile *t*, and the distance is measured in grid units (with adjacent tiles corresponding to one unit). Importantly, the directional nature implies *d_tq_* (*A*→*B*) # *d_qt_* (*A*→*B*) capturing asymmetric spatial relationships between cell types. The minimum directional distance from tile *t* of class *A* to class *B* was defined as:

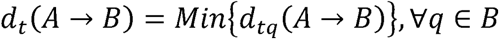

This procedure yielded a distribution of the minimum directional distances.

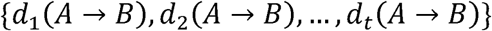

across all tiles of class *A*. For each sample, spatial distance summary statistics were computed based on the minimum directional distances between pairs of cell-type tiles. For every cell-type pair, we calculated the mean, median, standard deviation, and interquartile range (IQR) of these minimum weighted distances. The summary statistics are highly consistent with each other, and we choose to use median for downstream statistical modeling. Since normal epithelial cells were consistently underrepresented and did not support reliable distance estimation, it was excluded from spatial analyses.

Mutation burden was calculated from TCGA-COAD mutation annotation format files as the total number of somatic mutations per patient. Each median distance was regressed individually against log-transformed mutation burden, adjusted for age, gender, and cancer stage:

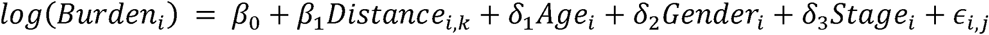

where *Distance_i,k_* represents median distance *k* of patient *i*, {*Age_i_*, *Gencter_i_*, *stage_i_*} are patient-level covariates, and ,*i,j* is the error term.

Following extraction of cell type proportions and directional spatial distance metrics, we conducted a series of statistical analyses to evaluate the clinical relevance of histopathological features in the TCGA-COAD cohort. Progression-free interval (PFI) was used as the primary clinical outcome. Patient survival data were obtained from the TCGA Clinical Data Resource supplemental tables, including PFI with corresponding time-to-event information. Cancer stage was standardized into four categories (Stage I–IV), with Stage I used as the reference. Continuous spatial distances (mean and median for each directed pairwise relationship) and overall cell type proportions were first examined through distributional assessments, including histograms and empirical cut-point estimation. To facilitate visualization and non-parametric comparison, overall cell type proportions were stratified into tertiles, age was grouped into quartiles, and median spatial distances were evaluated using binary median-split thresholds. Kaplan–Meier curves with log-rank tests were generated for all grouped variables.

Associations between each histopathological variable and survival outcomes were first assessed using marginal Cox proportional hazards models. Each model estimated the hazard ratio for a single feature, evaluated in its continuous form (distance summaries, proportions, clinical covariates) or binary form (median-split distances) if applicable:

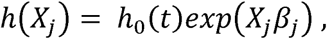

where *h*(*X_j_*) is the hazard function, *h*_0_ (*t*) is the baseline hazard, and *X_j_* denotes histopathological feature 1. To evaluate whether spatial metrics provided independent prognostic information, we additionally fitted multivariable Cox models adjusting for age, gender, and AJCC cancer stage. All hazard ratios, 95% confidence intervals, and Wald p-values were recorded. This analytical framework enabled systematic evaluation of both spatial architecture and compositional tumor ecology as predictors of patient outcomes in colorectal cancer.

## Data availability

The colorectal training data used in this study were collected from multiple sources. The single-cell RNA-seq reference data were obtained from the OSF repository (https://osf.io/hftq2/, DOI: 10.17605/OSF.IO/HFTQ2). Colorectal cancer H&E images and paired spatial transcriptomic data for training were collected from three independent cohorts. The first cohort was obtained from the Human Tumor Atlas Network Vanderbilt Atlas, available at the HTAN Data Coordinating Center portal (https://data.humantumoratlas.org/). The second cohort was obtained from the HEST-1K dataset developed by the Mahmood Lab (https://github.com/mahmoodlab/HEST). The third cohort consists of in-house colorectal cancer histopathology images generated by our group and the data are available from the corresponding authors upon reasonable request. The external colorectal cancer H&E images used for clinical association were downloaded from the National Cancer Institute Imaging Data Commons (IDC) portal (https://portal.imaging.datacommons.cancer.gov/) under the TCGA colon adenocarcinoma (TCGA-COAD) collection. The corresponding clinical data were obtained from the TCGA-CDR dataset (TCGA-CDR-SupplementalTableS1.xlsx), available through the Genomic Data Commons (GDC) data portal (https://api.gdc.cancer.gov/data/1b5f413e-a8d1-4d10-92eb-7c4ae739ed81).

The breast cancer training data were also collected from multiple public sources. The single-cell RNA-seq reference data were obtained from Mo et al. (PMID: 39478210) and are part of the HTAN dbGaP study accession phs002371.v3.p1, accessible through the HTAN Data Coordinating Center portal under the HTAN WUSTL Atlas (https://data.humantumoratlas.org/). Six breast cancer datasets were downloaded from 10x Genomics, including datasets from human breast cancer block A, whole-transcriptome breast cancer samples, ductal carcinoma in situ and invasive carcinoma FFPE samples, and fresh-frozen breast cancer tissues (https://www.10xgenomics.com/). Additional breast cancer spatial transcriptomics data were obtained from Wu et al. (PMID: 34493872), available through Zenodo (<u>10.5281/zenodo.4739739</u>); the corresponding full-resolution whole-slide images were obtained directly from the authors. This cohort included six samples. Further breast cancer data were obtained from Liu et al. (PMID: 36357424), available through GEO under accession GSE190811, including samples from four breast cancer patients. The external breast cancer H&E images used for clinical association were downloaded from the National Cancer Institute Imaging Data Commons (IDC) portal (https://portal.imaging.datacommons.cancer.gov/) under the TCGA breast invasive carcinoma (TCGA-BRCA) collection.

## Code availability

All code used for model development and data analysis in this study is publicly available on GitHub. The analysis pipelines for colorectal cancer and breast cancer are available at https://github.com/Sun-lab/STpath-CRC and https://github.com/Sun-lab/STpath-BC, respectively. The general STpath software package is available at https://github.com/Sun-lab/STpath-software.

## Acknowledgements

This work was supported by NIGMS R01 grant GM105785

## Author contributions

W.S. and Z.L. conceived the study. W.S. supervised the study. Z.S. initiated the methodological development of this work using convolutional neural network. S.C. subsequently adopted this framework and developed the computational methods using different foundation models, implemented the methods, performed the colorectal cancer analyses, generated the figures, and led the manuscript writing. Z.S. contributed to model implementation, breast cancer data analysis, and manuscript writing. K.A.M. provided pathology expertise and performed pathological evaluation to assess the biological consistency of model predictions. M.Y. and W.M.G. provided data resources and contributed to data interpretation. All authors reviewed and approved the final manuscript.

