## Supplementary Materials for "Translating Histopathology Foundation Model Embeddings into Cellular and Molecular Features for Clinical Studies"

### Supplementary Methods

#### Spatial Transcriptomics Data Integration

When creating image patches, we determine the appropriate patch radius by calculating the median distance between adjacent spots in 10X Visium spatial transcriptomic data. For each spatial transcriptomics matrix, we calculated the median distances between consecutive spots: $\tilde{d}_x$ represents the median horizontal distance between adjacent spots, and $\tilde{d}_y$ represents the median vertical distance. The radius $r$ was then calculated as:

$$r = \frac{\tilde{d}_x + \tilde{d}_y}{2}.$$

For each valid spot $s_i = (x_i, y_i)$ with $(in\_tissue = 1 \ \wedge \ tissue\_region \neq -1)$, a square patch was extracted centered at $s_i$ with side length $2r$. Patches extending beyond the image boundaries were excluded.

The detection of white patches was implemented using grayscale conversion with intensity threshold $\delta_{white} = 220$. The white ratio of a patch $P$ was defined as

$$white\_ratio(P) = \frac{|(x, y) \in P : gray(x, y) > \delta_{white}|}{|P|}$$

Patches that exceeded the cutoff point ($white\_ratio(P) \geq 0.4$) were removed. Regions with fewer than 30 remaining patches were excluded. These procedures were applied to both colorectal cancer and breast cancer data, excecpt HEST-1K data, which have already been preprocessed.

#### Single-cell RNA-seq Reference Data Processing (Colorectal Cancer)

We processed published colorectal cancer scRNA-seq data (Vanderbilt HTAN) containing pre-annotated cell types using the Python package Scanpy. To ensure annotation accuracy, we implemented a double-confirmation strategy combining original annotations with de novo clustering analysis.

Cells and genes were filtered according to the following criteria:

$$0.01 \times G_{total} \leq n_{genes}(c) \leq 2500,\ n_{cells}(g) \geq 0.01 \times C_{total},\ \%mt(c) < 5\%$$

where $G_{total}$ and $C_{total}$ are the total number of genes and cells of the scRNA-seq data, $n_{genes}(c)$ is the number of expressed genes per cell $c$, $n_{cells}(g)$ is the number of cells expressing gene $g$, and $\%mt(c)$ is the mitochondrial gene percentage per cell. Normalization and transformation steps included:

$$x_{c,g}^{norm} = \log\left(\frac{10^4 \cdot x_{c,g}}{\sum_j x_{c,g}} + 1\right),$$

where $x_{c,g}$ is the raw count of gene $g$ in cell $c$. PCA was computed with the first 40 principal components. A $k$-nearest neighbor graph was constructed ($k = 10$) using the top 40 PCs. Leiden clustering (resolution = 0.5) produced de novo cell clusters. UMAP was applied for visualization. De novo clusters were then cross-referenced with original annotations to confirm final cell type identities. The double confirmation means that we assign a cell to a cell type $T$ only if it was annotated to cell type $T$ by the original annotation and it belongs to the cluster dominated by cell type $T$.

After double confirmation, marker genes were identified using the Wilcoxon Rank-Sum test with Benjamini-Hochberg correction for FDR control. We conducted seven types of pairwise comparisons:
(1). Each cell type vs. all other groups.
(2). Individual cancer subtypes (ASC I-III, CSC I-IV, SSC) vs. normal epithelial group (CT, EE, TUF, ABS).
(3). Individual normal epithelial subtypes (CT, EE, TUF, ABS) vs. cancer group (ASC I-III, CSC I-IV, SSC).
(4). CD4+ T vs. CD8+ T cells.
(5). CD8+ T vs. CD4+ T cells.
(6). FIB vs. END.
(7). END vs. FIB.
A gene $g$ was selected as a marker for a cell type group $t$ if:

$$p_{adj}(g) < 0.05, \quad \log_2 FC > 1.0, \quad f_t(g) > 0.25$$

where $p_{adj}(g)$ is the adjusted p-value, $\log_2 FC$ is the log2 fold change between a target cell type group and the comparison group, and $f_t(g)$ is the expression frequency of the gene $g$ in the cell type group $t$. From comparison (1), the 50 best genes ranked by $\log_2 FC$ were selected for each cell type. From comparisons (2) - (7), the top 40 genes were selected for each corresponding subtype. This resulted in 899 total marker genes after merging all selected genes.

#### Single-cell RNA-seq Reference Data Processing (Breast Cancer)

Publicly available single-cell RNA sequencing (scRNA-seq) datasets from Wu et al. (2021) and Mo et al. (2024) (details are provided in the Data Availability section of the main text) were obtained to construct a comprehensive reference for breast cancer. Raw gene expression matrices were processed using the Seurat R package. To ensure annotation accuracy, we implemented a double-confirmation strategy combining original author annotations with de novo integrated clustering analysis.

Standard quality control was performed for each source. Cells and genes were filtered according to the following criteria:

$$200 \leq n_{genes}(c) \leq 6000,\ n_{cells}(g) \geq 3,\ \%mt(c) < 15\%$$

where $n_{genes}(c)$ is the number of detected genes per cell $c$, $n_{cells}(g)$ is the number of cells expressing gene $g$, and $\%mt(c)$ is the mitochondrial gene percentage per cell.

To define cell populations robustly across datasets, normalization and variance stabilization were performed using `SCTransform`, followed by integration. Dimensionality reduction was computed with the first 30 principal components. A $k$-nearest neighbor graph was constructed using these PCs. Louvain clustering (resolution = 0.5) produced de novo integrated clusters. UMAP was applied for visualization.

We utilized a "divide-and-conquer" iterative purification strategy. Initial clusters were broadly annotated using canonical marker genes (e.g., CD3D for T cells, MS4A1 for B cells, EPCAM for epithelial cells). Clusters exhibiting ambiguous or mixed signatures were isolated and re-clustered at high resolution to resolve distinct subpopulations. We applied a strict consensus filtering approach to generate "pure" reference datasets. For each cell, the de novo integrated cluster assignment was cross-referenced with the original authors' major cell type annotations to confirm final cell type identities. We assign a cell to a cell type $T$ only if it was annotated to cell type $T$ by the original study and it belongs to the integrated cluster dominated by cell type $T$. This process resulted in a curated reference consisting of nine core cell types: T cells, B cells, Plasma cells, Plasmablasts, Myeloid cells, Cancer Epithelial, Normal Epithelial, Endothelial cells, and a combined Cancer-Associated Fibroblast (CAF)/Perivascular-like (PVL) group.

After double confirmation, we constructed a robust consensus gene signature derived from both reference datasets. Differential expression analysis (Wilcoxon rank-sum test) was performed independently on the purified Wu et al. and Mo et al. datasets to identify top marker genes for each cell type. A gene $g$ was selected as a marker for a cell type group $t$ if:

$$p_{adj}(g) < 0.05, \quad \log_2 FC > 1.0, \quad f_t(g) > 0.25$$

where $p_{adj}(g)$ is the Bonferroni-adjusted p-value, $\log_2 FC$ is the log2 fold change between the target cell type group and the background, and $f_t(g)$ is the expression frequency of gene $g$ in the cell type group $t$.

We defined the final signature gene list by taking the intersection of the top differentially expressed genes from both datasets for each cell type, ensuring that only markers consistently upregulated across studies were retained. This data-driven list was supplemented with a curated panel of established breast cancer cell type markers obtained provided by Wu et al. and Mo et al.

The final reference for CARD was generated by extracting the raw single-cell count matrices from the purified Seurat objects, strictly subsetted to this consensus signature gene list, thereby prioritizing biological signal over dataset-specific variation.

#### Deconvoluted Results Validation

Cell type-specific marker genes were identified from the same scRNA-seq reference dataset that was used for CARD deconvolution, employing differential expression analysis. Gene expression counts were first normalized by library size to account for sequencing depth differences:

$$X_{c,g}^{CN} = \frac{X_{c,g}}{\sum_g X_{c,g}}$$

where $X_{c,g}$ is the raw count of cell $c$ and gene $g$. Then for a target cell type $t$. we calculated the average of target cell type as well as the non-target:

$$\bar{X}_{t,g}^{CN} = \frac{\sum_{i \in t} X_{c,g}^{CN}}{N_t}, \quad \bar{X}_{\sim t,g}^{CN} = \frac{\sum_{c \notin t} X_{c,g}^{CN}}{N_{\sim t}}$$

where $N_t$ is the number of single cells of the cell type $t$, $N_{\sim t}$ is the number of single cells of all other cell types except for $t$. For each target cell type $t$ and each gene $g$, we calculated the $\log_2 FC$ between target cell type expression and all other cell types:

$$\log_2 FC_{t,g} = \log_2(\bar{X}_{t,g}^{CN} + 1 \times 10^{-8}) - \log_2(\bar{X}_{\sim t,g}^{CN} + 1 \times 10^{-8})$$

Genes with $\log_2 FC_{t,g} \geq 1$ were retained as candidate markers for cell type $t$, and ranked by expression specificity. We grouped the 20 fine-grained cell types into 5 major categories: Tumor Cells (8 subtypes: ASC I-III, CSC I-IV, SSC I), Normal Epithelial Cells (4 subtypes: ABS, CT, EE and TUF), T Cells (2 subtypes: CD4+T and CD8+T), Other Immune Cells (4 subtypes: PLA, MYE, MAS and B), and Stromal Cells (2 subtypes: FIB and END). For each major cell type category, we selected marker genes using a tiered approach: the top 5 marker genes from each of the 8 cancer cell subtypes were combined to create the Cancer Cells signature, the top 10 marker genes from each of the 4 normal epithelial cell subtypes formed the Normal Epithelial Cells signature, the top 40 marker genes were selected for T Cells, the top 10 marker genes from each of the 4 immune cell subtypes comprised the Other Immune Cells signature, and the top 20 marker genes from fibroblasts and endothelial cells each were combined for the Stromal Cells signature. So, the 20 cell types $t$ were grouped into 5 big cell type categories $T$.

For each spatial spot, we calculated the relative marker gene expression (RMGE) as a proxy for true cell type abundance. To account for baseline expression differences across cell types, the count normalized expression was further divided by the average expression of each marker gene in the corresponding cell type from the CARD signature matrix (B-matrix).

$$\bar{X}_{T,g}^{CN} = \frac{\sum_{i \in T} X_{i,g}^{CN}}{N_T}, \quad \bar{X}_{T,g}^{CN\_BN} = \frac{\bar{X}_{T,g}^{CN}}{\sum_{t \in T} B_{t,g}/|T|}, \ \forall \ g \in G_T$$

$$RMGE_T = \frac{\sum_{g \in G_T} \bar{X}_{T,g}^{CN\_BN}}{|G_T|}$$

where $\bar{X}_{T,g}^{CN}$ is the averaged count normalized gene expression of gene $g$ and cell type $T$, $\bar{X}_{T,g}^{CN\_BN}$ is the B-matrix normalized version of $\bar{X}_{T,g}^{CN}$, $N_T$ is the total number of cells of cell type $T$, $G_T$ is the marker genes list of cell type $T$, $|T| = 5$ is the total number of big cell type categories. The relative expression values for all marker genes within each cell type category were then averaged and converted to proportions by normalizing across all five cell type categories to sum to 1.

$$RMGE_T^{Norm} = \frac{RMGE_T}{\sum_{T'} RMGE_{T'}}$$

The validation principle relies on the biological expectation that deconvoluted cell type proportions should correlate with marker gene-based evidence from the same tissue. For each tissue region, we performed Spearman correlation analysis between deconvoluted cell type proportions (DCTP) and normalized RMGE. The correlation was assessed using Spearman's correlation coefficients ($\rho$) indicating the strength of monotonic relationships. For each cell type and tissue region, we recorded sample size (number of spatial spots), and Spearman correlation coefficient. For each grouped cell type $T$, We calculate the weighted average of Spearman's correlation using sample size as the weights, so that we get $\bar{\rho}_T$. and any samples with Spearman's correlation $\geq 1.1 \times \bar{\rho}_T$ were selected as our final samples for downstream models training.

#### The Complementary analysis of Foundation Models

We quantified each foundation model's contribution to the combined model using a comprehensive feature importance decomposition approach. In the combined models, features from the five foundation models were concatenated in a fixed order: UNI2h features (indices 0-460), Virchow features (indices 461-1228), Virchow2 features (indices 1229-1996), ProvGigapath features (indices 1997-2457), and Conch features (indices 2458-2611). For each cell type XGBoost models, we calculated two metrics: Absolute Contribution and Normalized Contribution.

To calculate Absolute Contribution, we first computed the mean feature importance across all cross-validation folds for the combined XGBoost model. Then, for each foundation model $m$, the absolute contribution was calculated as:

$$C_{abs}^{m} = \sum_{k \in \mathcal{F}(m)} I_k$$

where $\mathcal{F}(m)$ represents the set of feature indices belonging to model $m$, and $I_k$ is the importance score of feature $k$. The percentage contribution was then computed as:

$$P_{abs}^{m} = \frac{C_{abs}^{m}}{\sum_{m'} C_{abs}^{m'}} \times 100\%$$

To account for the varying number of features contributed by each model, we calculated the Normalized Contribution as:

$$C_{norm}^{m} = \frac{C_{abs}^{m}}{|\mathcal{F}(m)|}$$

where $|\mathcal{F}(m)|$ is the number of features from model $m$. The normalized percentage contribution was:

$$P^m_{norm} = \frac{C^m_{norm}}{\sum_{m'} C^{m'}_{norm}} \times 100\%$$

This dual-metric approach revealed both the raw predictive power of each model (absolute contribution) and the per-feature efficiency (normalized contribution). The analysis demonstrated model-specific strengths: some models contributed many moderately important features, while others provided fewer but highly discriminative features, indicating complementary roles in capturing different aspects of tissue morphology for each cell type.

To evaluate the complementarity of feature representations across foundation models, we conducted pairwise Ridge regression analyses. For each model pair, the standardized features of one model were regressed against those of the other, and the coefficient of determination ($R^2$) was computed. Lower $R^2$ values reflected greater complementarity, whereas higher values suggested redundancy between models.

Formally, for each pair of foundation models $m_i$ and $m_j$, the standardized feature matrices were modeled as:

$$\boldsymbol{Y}^{(m_i)} = \boldsymbol{X}^{(m_j)}\boldsymbol{\beta} + \boldsymbol{\epsilon}$$

where $\boldsymbol{Y}^{(m_i)}$ denotes the standardized features from model $m_i$, $\boldsymbol{X}^{(m_j)}$ denotes those from model $m_j$, $\boldsymbol{\beta}$ are the ridge regression coefficients, and $\boldsymbol{\epsilon}$ is the residual term.

The ridge regression objective with $L_2$ regularization was defined as:

$$\min_{\boldsymbol{\beta}} ||\boldsymbol{Y}^{(m_i)} - \boldsymbol{X}^{(m_j)}\boldsymbol{\beta}||_2^2 + \alpha||\boldsymbol{\beta}||_2^2$$

with the regularization parameter fixed at $\alpha = 1.0$

$$R^2_{m_i \leftarrow m_j} = 1 - \frac{\sum(\boldsymbol{Y}^{(m_i)} - \hat{\boldsymbol{Y}}^{(m_i)})^2}{\sum(\boldsymbol{Y}^{(m_i)} - \bar{\boldsymbol{Y}}^{(m_i)})^2}$$

where $\hat{\boldsymbol{Y}}^{(m_i)} = \boldsymbol{X}^{(m_j)}\hat{\boldsymbol{\beta}}$ are the predicted features and $\bar{\boldsymbol{Y}}^{(m_i)}$ is the mean of the target features.

This analysis revealed that low $R^2$ values (approaching zero) indicated limited predictive alignment between models, thereby highlighting strong complementarity. In contrast, higher $R^2$ values reflected overlapping or redundant feature representations.

### Supplementary Figures

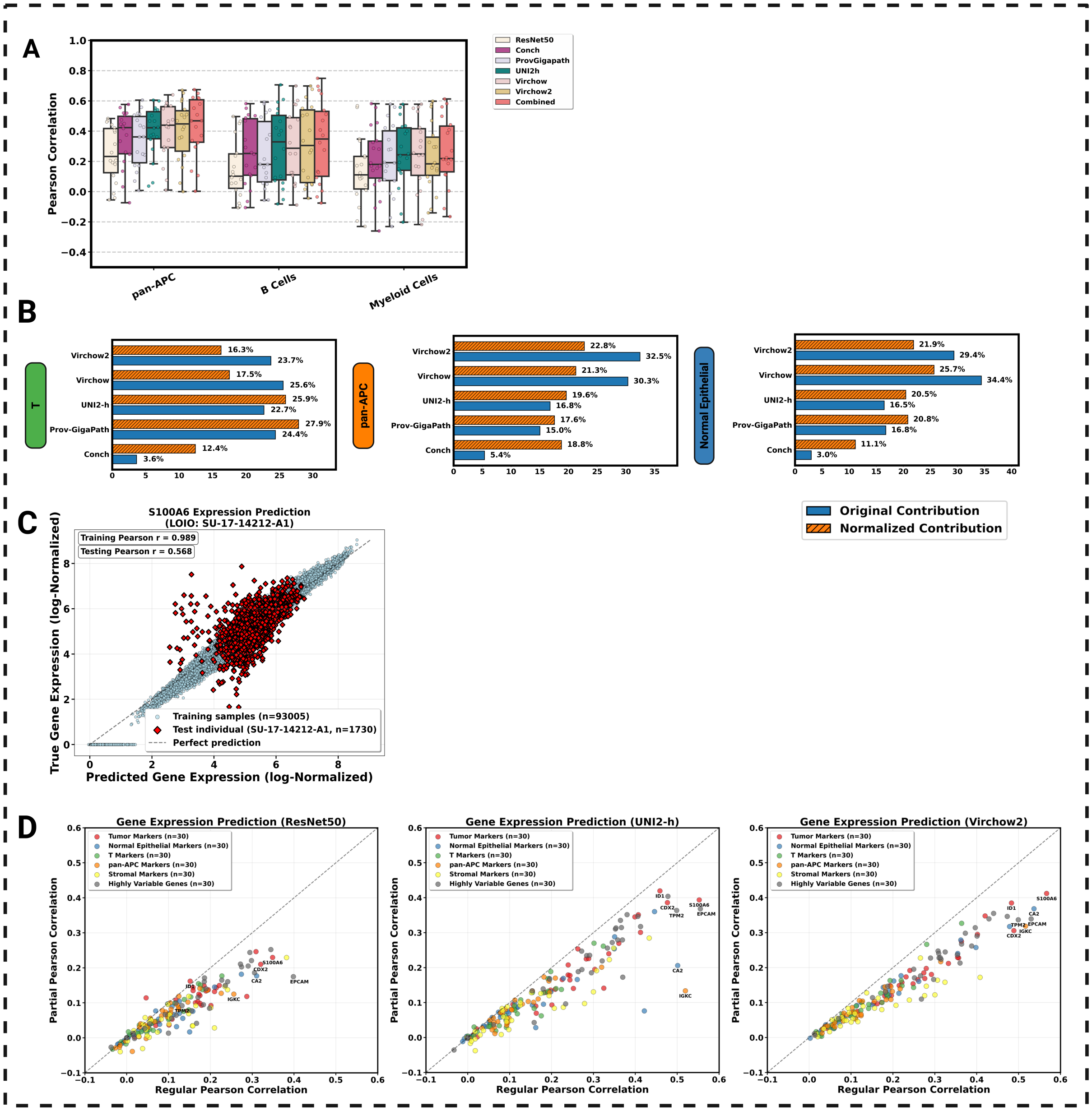

**Figure S1. pan-Antigen Presenting cells (pan-APCs) prediction analysis, model contribution analysis, example gene expression prediction, and partial correlation analysis.** (A). Comparison of prediction accuracy of B cells, Myeloid cells, or combining them to one group of pan-APCs. (B). Contribution analysis of foundation models to the combined XGBoost prediction performance for T, pan-APC and normal epithelial cells. (C). Example of **S100A6** expression prediction in one held-out individual (SU-17-14212-A1) under LOIO cross-validation using Virchow2. (D). Comparison of regular Pearson correlation versus partial Pearson correlation controlling for cell type proportions across 180 predictor genes for three representative models. Genes include 150 cell type-specific markers (30 each for Tumor, Normal epithelial, T cells, Stromal, and pan-APCs) and 30 highly variable non-marker genes. Each dot represents one gene. Seven representative genes are highlighted: five known cell type markers (ID1, CDX2, S100A6, CA2, IGKC) and two highly variable genes (EPCAM, TMP2).

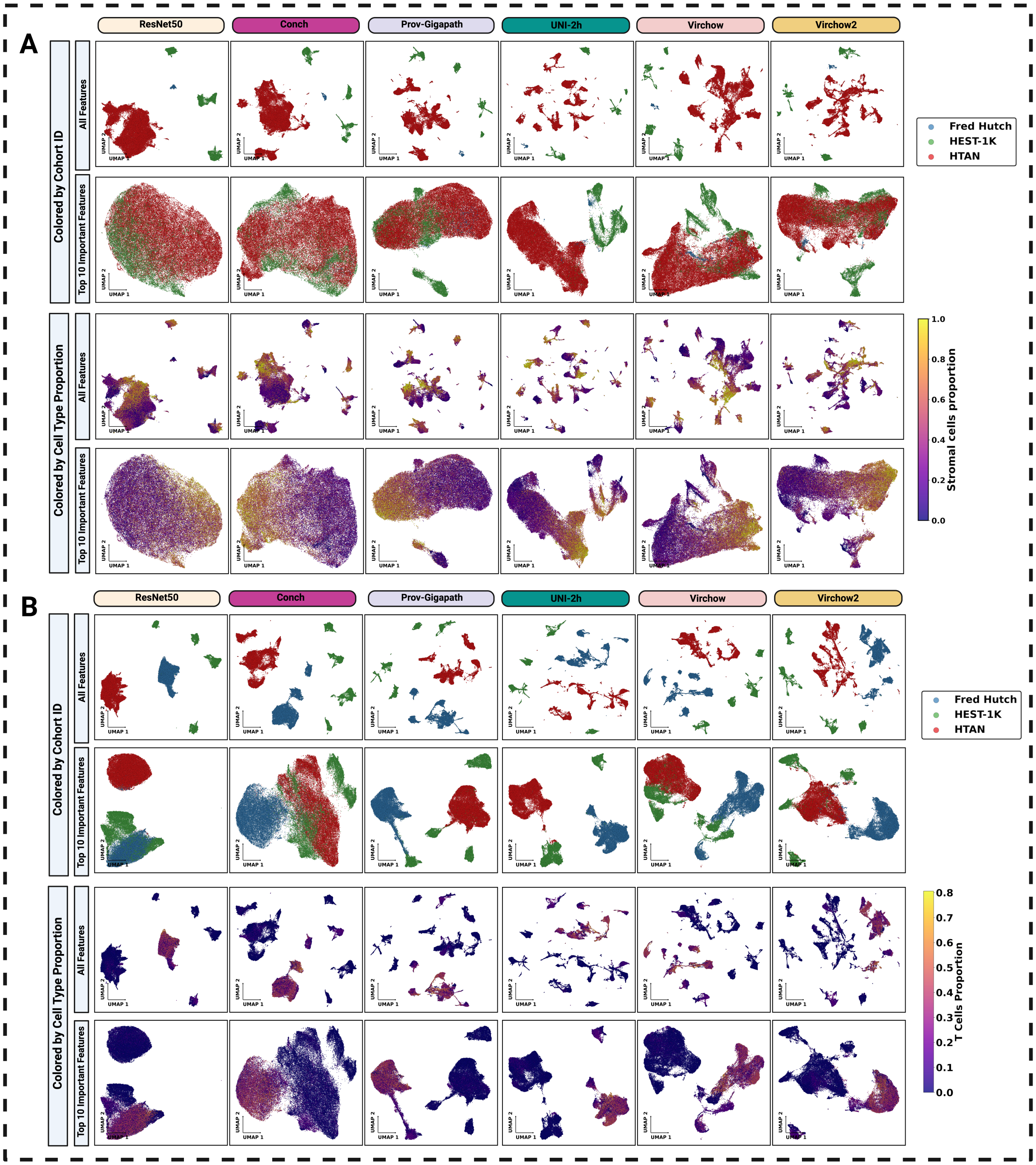


**Figure S2. UMAP visualizations of the image tiles used to train stromal and T cells proportion models.** (A). Embeddings are shown for the full feature set and for the top 10 XGBoost selected features (rows). Points are colored by cohorts (top panels) and by deconvolved stromal cell proportion (bottom panels; color bar at right). (B). Same layout as (A), but for T cells.

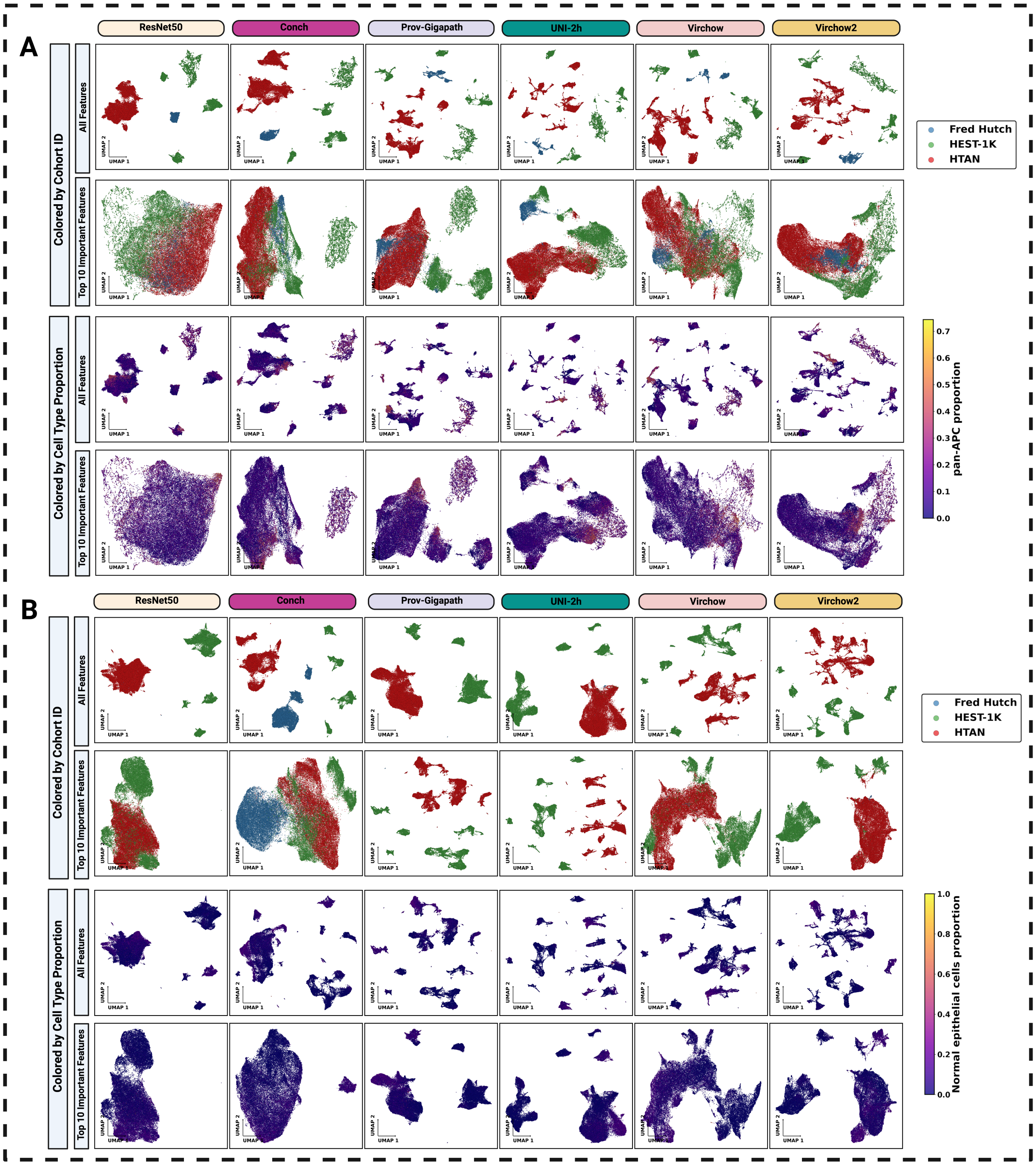


**Figure S3. UMAP visualizations of the image tiles used to train pan-APC and normal epithelial cells proportion models.** (A). Embeddings are shown for the full feature set and for the top 10 XGBoost selected features (rows). Points are colored by cohorts (top panels) and by deconvolved pan-APC proportion (bottom panels; color bar at right). (B). Same layout as (A), but for normal epithelial cells.

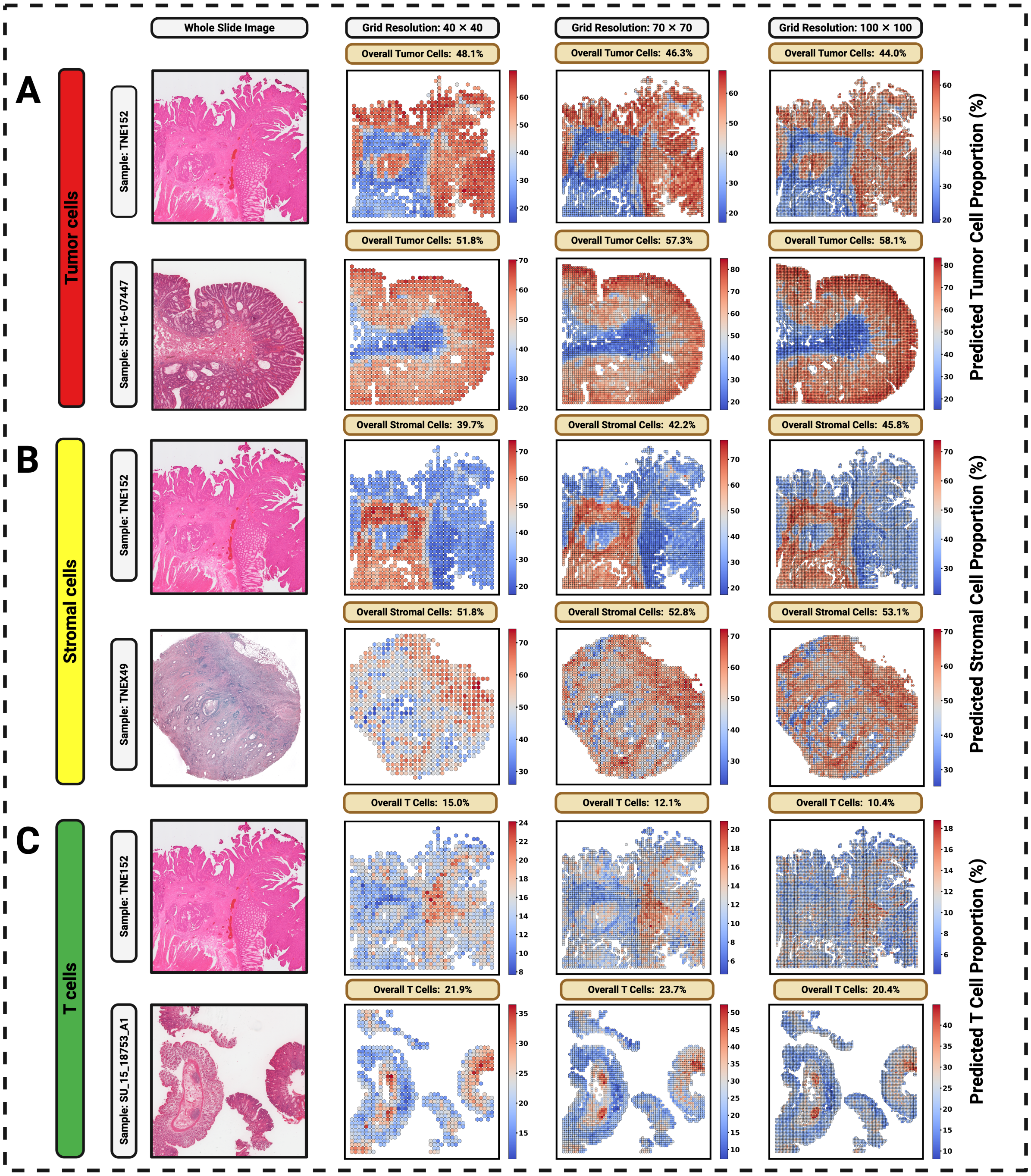


**Figure S4. Grid-resolution robustness of predicted cell-type proportion maps.** (A). Tumor-enriched WSIs: original image and hexagonal binned heatmaps at grid resolutions 40×40, 70×70, and 100×100; slide-level overall tumor proportions are annotated above each map. (B). Stromal-enriched WSIs shown at the same three grid resolutions with overall stromal proportions annotated. (C). T-cell–enriched WSIs shown at the same three grid resolutions with overall T-cell proportions annotated.

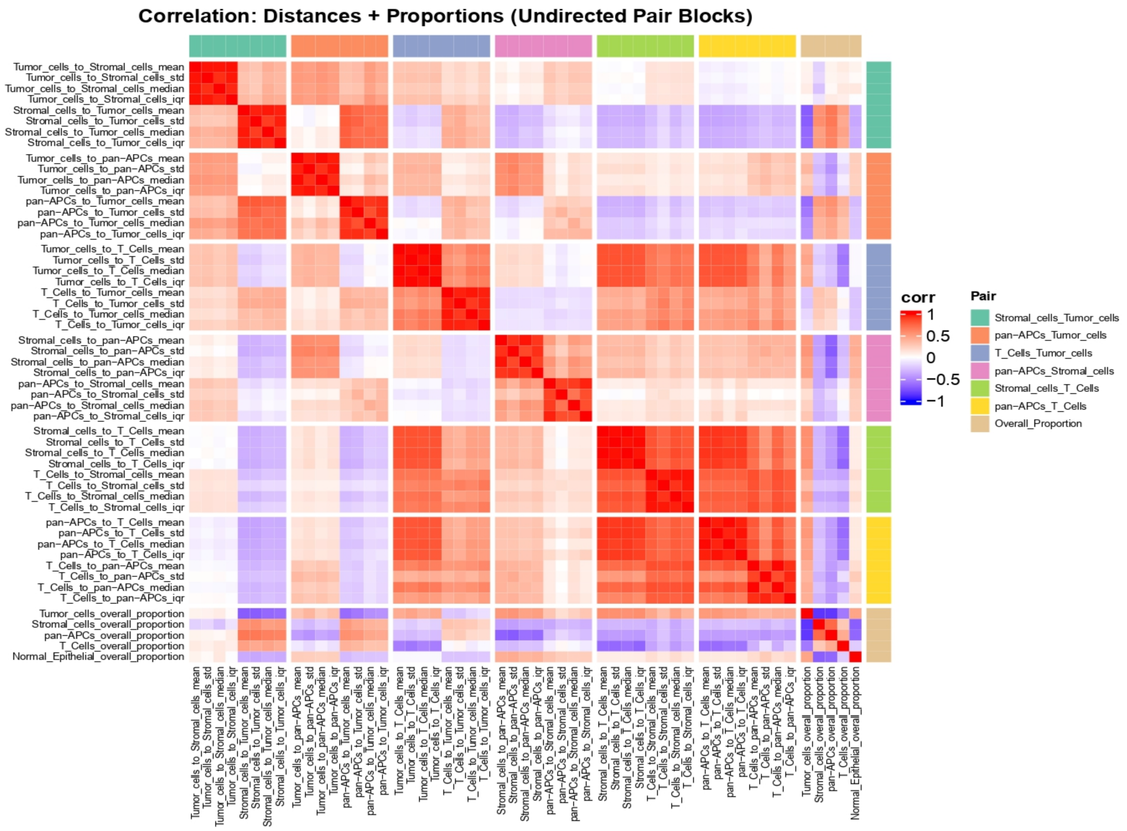


**Figure S5. Correlation heatmap showing relationships between different summary statistics of minimum directional distance.** Pearson correlation coefficients were computed among four summary statistics (mean, std, median, IQR) for six cell-type pairs and overall cell-type proportions across TCGA COAD samples.

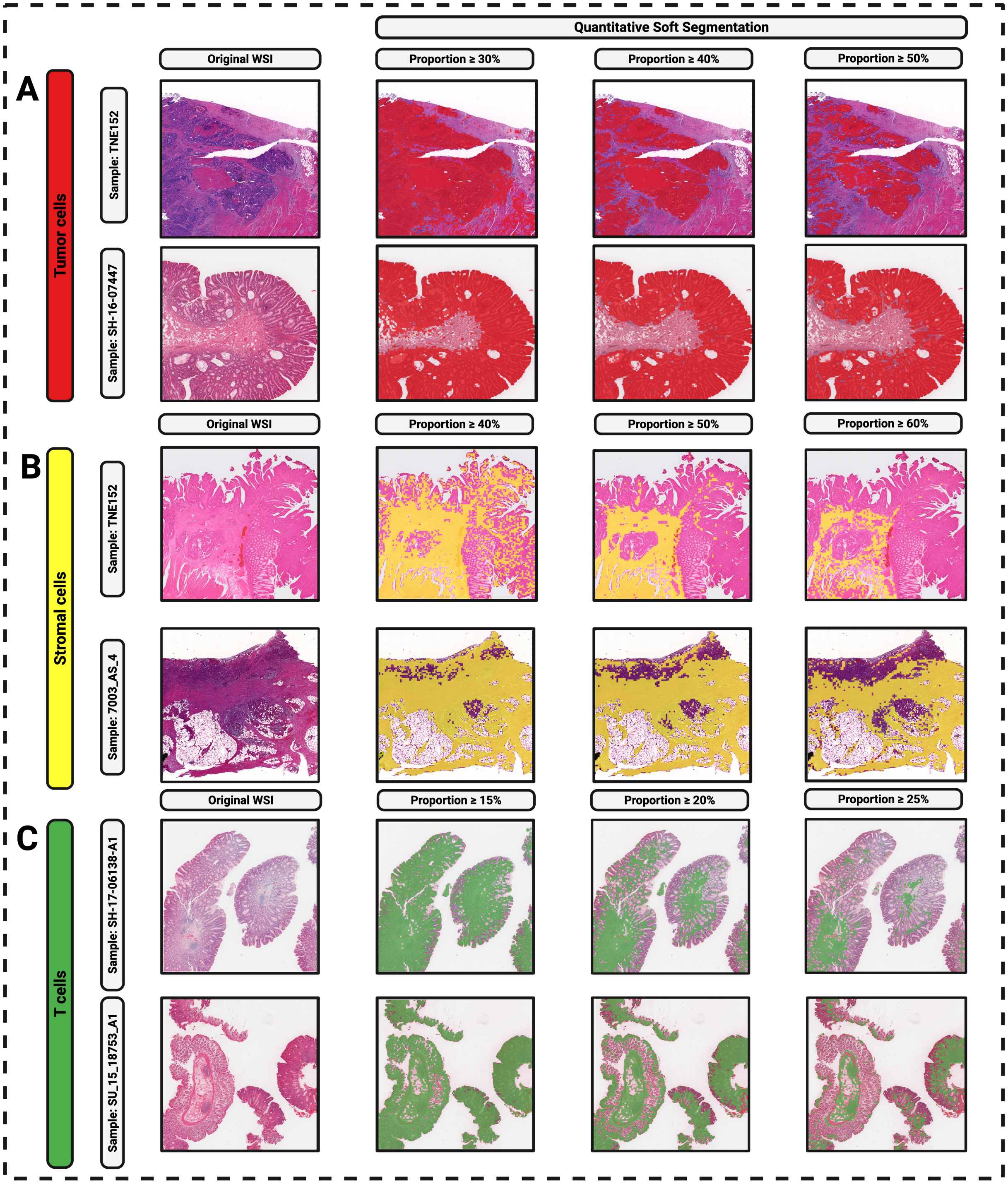


**FIgure S6. Quantitative soft tissue segmentation from predicted cell-type proportions.** (A). Tumor-enriched samples. For each WSI, the original image (left) is shown alongside transparent overlays highlighting tiles with predicted tumor proportions $\geq 30\%$, $\geq 40\%$, and $\geq 50\%$. (B). Stromal-enriched samples with overlays at thresholds $\geq 40\%$, $\geq 50\%$, and $\geq 60\%$. (C). T-cell–enriched samples with overlays at thresholds $\geq 15\%$, $\geq 20\%$, and $\geq 25\%$.

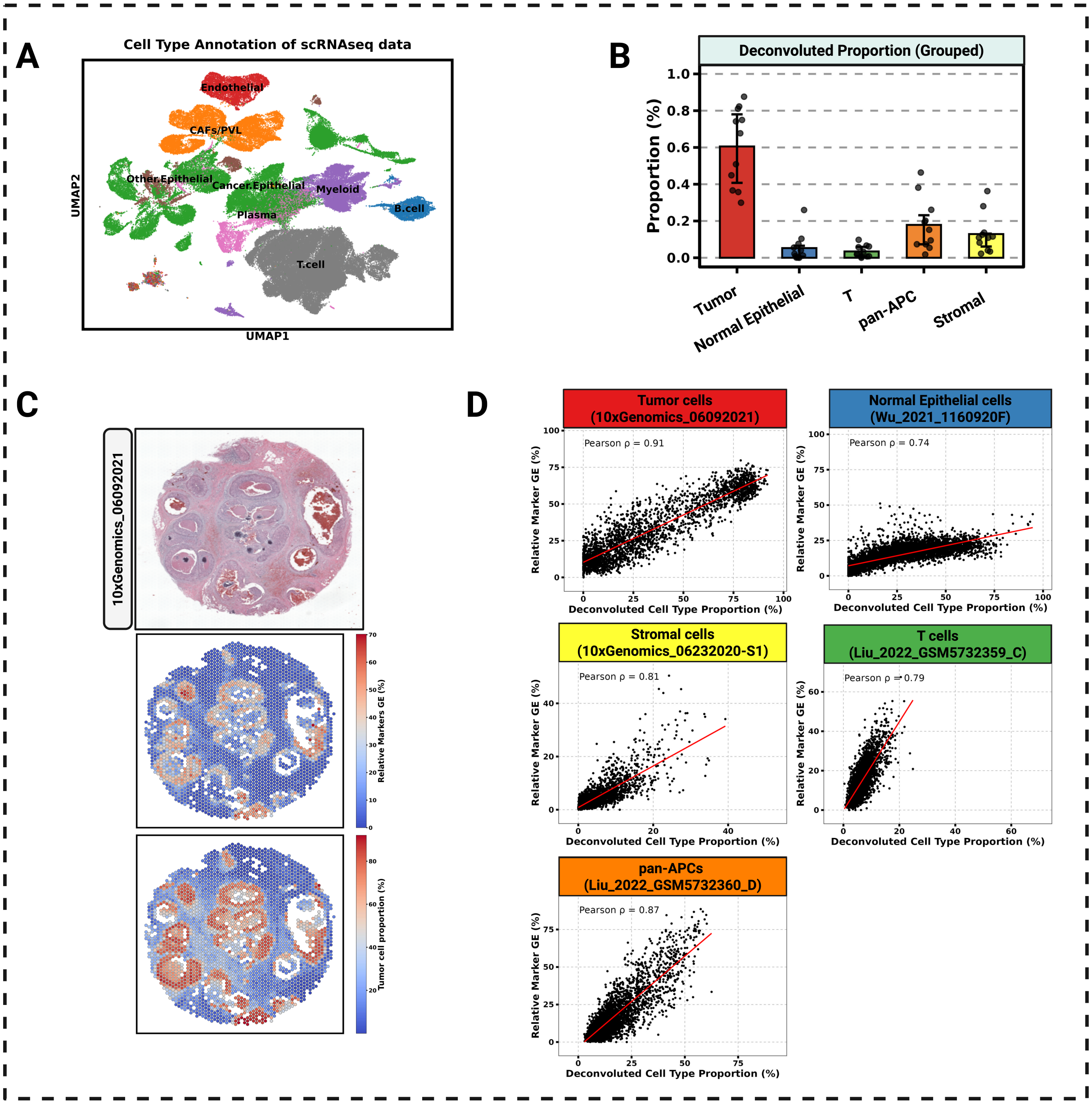


**Figure S7. CARD deconvolution and validation for breast cancer spatial transcriptomics data.** (A). UMAP visualization of scRNA-seq reference dataset showing eight major breast cancer cell populations: cancer epithelial, other epithelial, CAFs/PVL, myeloid, B cells, plasma cells, T cells, and endothelial cells. (B). Distribution of deconvoluted cell type proportions (grouped) across breast cancer spatial transcriptomics samples. Bars show median proportions with interquartile ranges. (C). Spatial distribution of cancer epithelial cells in a representative breast cancer sample. Top: H&E-stained section. Middle: Relative marker gene expression. Bottom: CARD-deconvoluted tumor proportion. (D). Validation of deconvolution accuracy. Scatter plots comparing deconvoluted cell type proportions versus relative marker gene expression for all eight cell types. Pearson correlation coefficients ($r = 0.74 - 0.91$) confirm accurate deconvolution.

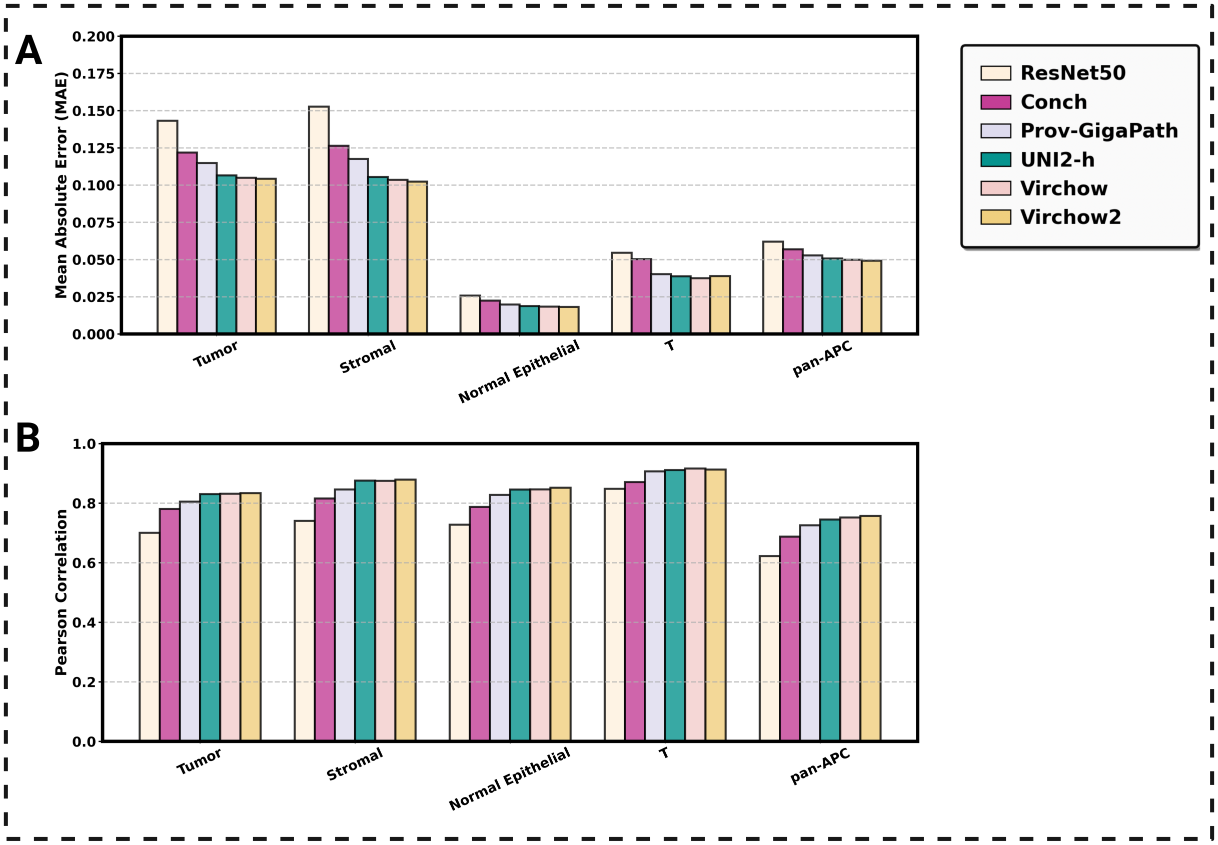


**Figure S8. Naive cross-validation performance.** (A). Mean Absolute Error (MAE) comparison across six models for five cell types using pooled 80% training and 20% testing split of all tiles. (B). Pearson correlation coefficients between predicted and true cell type proportions for the same models and cell types.

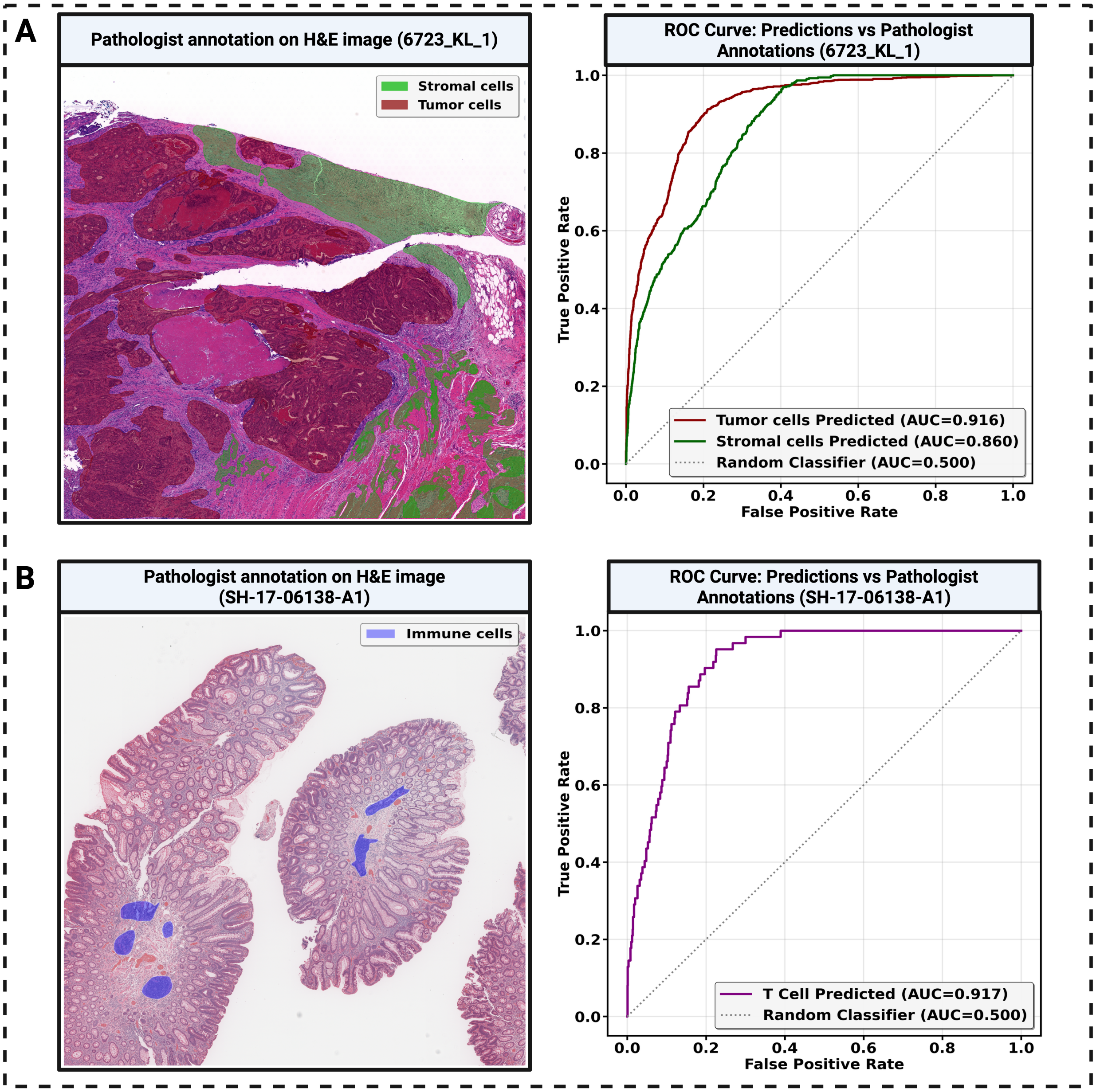


**Figure S9. Validation of cell type predictions using pathologist annotations in colorectal cancer samples.** (A) Left: Pathologist annotated regions for tumor cells (red) and stromal cells (green) overlaid on H&E image from sample 6723_KL_1. Right: Performance evaluation of predicted tumor and stromal cell proportions. The pathologist-annotated regions were used as ground truth (inside annotation = 1, outside = 0), and predictions were generated using models trained with this sample excluded from the training set. (B) Left: Pathologist-annotated immune cell regions (blue) on H&E image from sample SH-17-06138-A1. Right: Performance evaluation of predicted T cell proportions against pathologist-annotated immune regions.

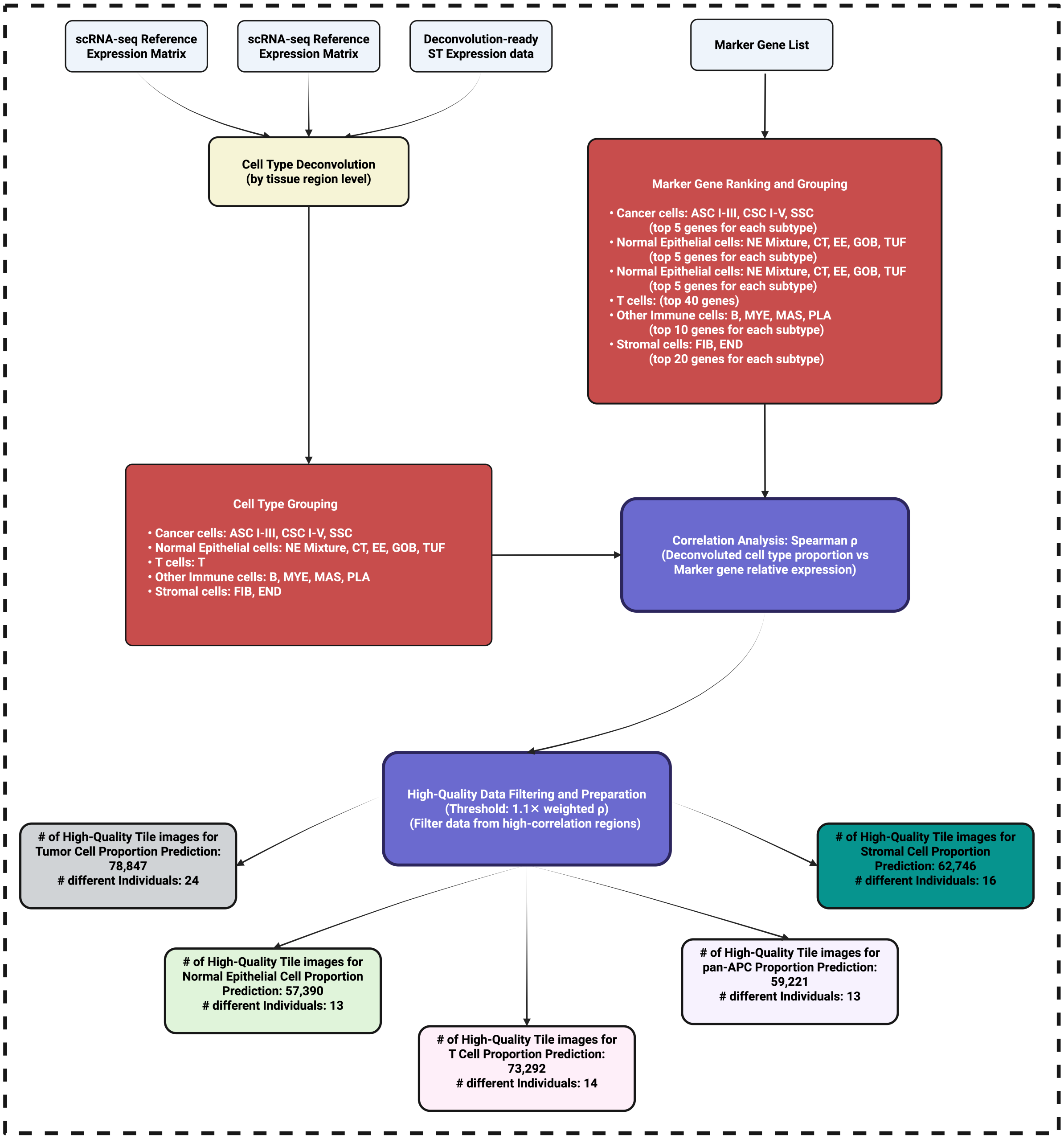


**Figure S10. Aligning H&E WSIs with spatial transcriptomics and generating patch-level inputs for cell type deconvolution.** Raw inputs include whole slide H&E images and ST spot coordinates together with the ST expression matrix. WSIs are aligned to ST coordinates via affine transformation, followed by tissue-region segmentation. Small regions with insufficient spots are removed to retain valid regions. Image tiles are then generated by determining the patch radius and removing over-white and corner tiles. Finally, tiles are organized and merged with the ST expression by region to produce the two outputs: (1) deconvolution-ready ST expression files and (2) matched image tiles.

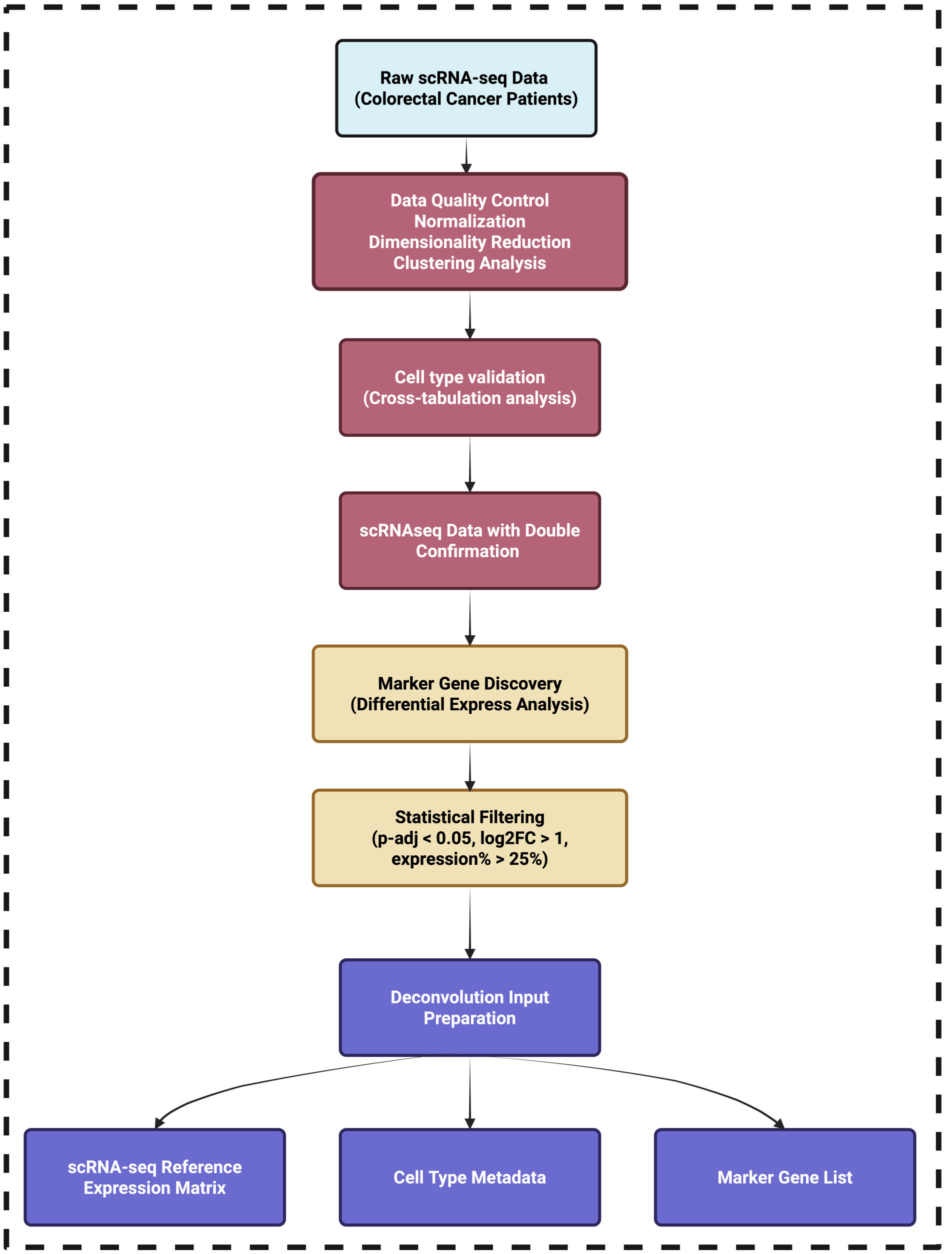


**Figure S11. Aligning H&E WSIs with spatial transcriptomics and generating patch-level inputs for cell type deconvolution.** Workflow illustrating the processing of raw scRNA-seq data from colorectal cancer patients. Following quality control, normalization, dimensionality reduction, and clustering, cell types were validated through cross-tabulation analysis. Marker genes were identified via differential expression analysis and subjected to statistical filtering (adjusted $p < 0.05$, $log2(F\_C) > 1$, expression$\% > 25\%$). The resulting curated dataset was used to prepare the reference expression matrix, cell type metadata, and marker gene list for subsequent cell type deconvolution.

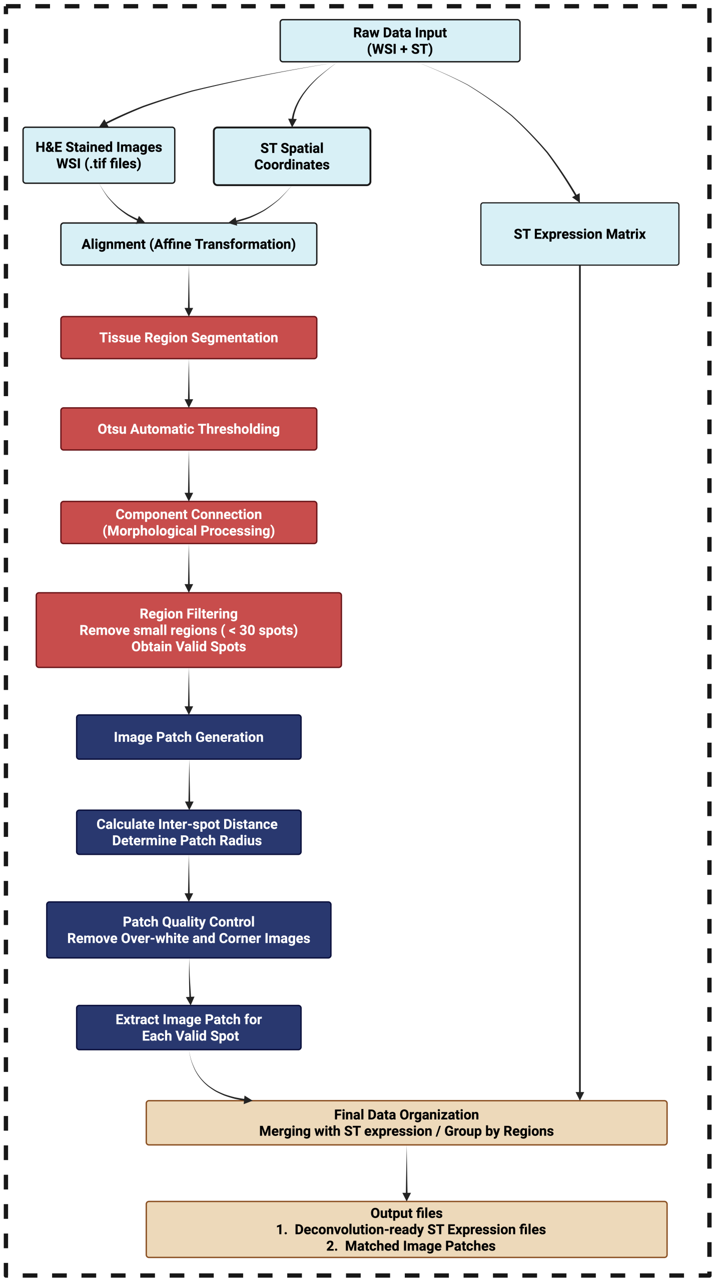


**Figure S12. Workflow for cell type deconvolution, validation of cell type deconvolution and preparation of high-quality tile images (Colorectal cancer).** Cell type deconvolution was first performed at the tissue region level, followed by cell type grouping into five major categories: Cancer cells, Normal epithelial cells, T cells, pan-APCs, and Stromal cells. Marker genes were ranked and selected for each subtype, and correlation analysis (Spearman's $\rho$) was used to validate deconvoluted cell type proportions against marker gene expression. High-quality data were then filtered based on correlation thresholds, yielding tile images with matched cell type proportions for downstream predictive modeling.

### Supplementary Tables

#### Table S1. Training samples and tile counts (Colorectal cancer).

| Sample ID | Source | Training for | Total Tiles |
|---|---|---|---|
| 6723_KL_1 | Vanderbilt HTAN | Tumor cell/ Normal epithelial cell/ Stromal cell | 3640 |
| 6723_KL_2 | Vanderbilt HTAN | Tumor cell / Normal epithelial cell/ Stromal cell/ T cell/ pan-APC | 3999 |
| 6723_KL_4 | Vanderbilt HTAN | T cell | 3355 |
| 7003_AS_1 | Vanderbilt HTAN | Normal epithelial cell/ T cell | 320 |
| 7003_AS_2 | Vanderbilt HTAN | Normal epithelial cell/ T cell | 117 |
| 7003_AS_3 | Vanderbilt HTAN | Normal epithelial cell/ Stromal cell | 2205 |
| 7003_AS_4 | Vanderbilt HTAN | Normal epithelial cell/ Stromal cell/ pan-APC | 2734 |
| 7003_AS_5 | Vanderbilt HTAN | Tumor cell/ Stromal cell/ T cell | 3052 |
| 7003_AS_6 | Vanderbilt HTAN | Tumor cell/ Stromal cell/ T / pan-APC | 2349 |
| 7003_AS_7 | Vanderbilt HTAN | Tumor cell/ Normal epithelial cell/ Stromal cell/ T cell/ pan-APC | 3216 |
| 7003_AS_8 | Vanderbilt HTAN | Tumor cell/ Normal epithelial cell/ pan-APC | 3798 |
| 7319_AS_2 | Vanderbilt HTAN | Stromal cell/ T cell | 1449 |
| 7319_AS_3 | Vanderbilt HTAN | Normal epithelial cell/ T cell/ pan-APC | 274 |
| 7794_AS_1 | Vanderbilt HTAN | Tumor cell/ Stromal cell/ T cell/ pan-APC | 638 |
| 7794_AS_2 | Vanderbilt HTAN | Tumor cell/ Normal epithelial cell/ Stromal cell/ T cell/ pan-APC | 572 |
| 8270_AS_1 | Vanderbilt HTAN | T cell/ pan-APC | 623 |
| 8270_AS_10 | Vanderbilt HTAN | Tumor cell/ Normal epithelial cell/ Stromal cell | 3698 |
| 8270_AS_12 | Vanderbilt HTAN | T cell/ pan-APC | 3268 |
| 8270_AS_2 | Vanderbilt HTAN | T cell | 562 |
| 8270_AS_5 | Vanderbilt HTAN | Tumor cell/ T cell/ pan-APC | 74 |
| 8270_AS_6 | Vanderbilt HTAN | Tumor cell/ Normal epithelial cell/ Stromal cell | 2623 |
| 8270_AS_7 | Vanderbilt HTAN | Tumor cell/ Stromal cell/ pan-APC | 3747 |

**Table S1 continued**

| Sample ID | Source | Training for | Total Tiles |
|---|---|---|---|
| 8270_AS_8 | Vanderbilt HTAN | Tumor cell/ Normal epithelial cell/ Stromal cell | 4106 |
| 8270_AS_9 | Vanderbilt HTAN | Tumor cell/ Stromal cell/ T cell | 3180 |
| 8578_AS_1 | Vanderbilt HTAN | Tumor cell/ Normal epithelial cell/ Stromal cell/ T cell/ pan-APC | 527 |
| 8578_AS_2 | Vanderbilt HTAN | Tumor cell/ Normal epithelial cell/ Stromal cell/ T cell/ pan-APC | 603 |
| 8578_AS_3 | Vanderbilt HTAN | Tumor cell/ Normal epithelial cell/ Stromal cell/ T cell/ pan-APC | 396 |
| 8899_AS_1 | Vanderbilt HTAN | Tumor cell/ Stromal cell/ pan-APC | 3652 |
| 8899_AS_3 | Vanderbilt HTAN | Tumor cell/ pan-APC | 1576 |
| 8899_AS_5 | Vanderbilt HTAN | Tumor cell/ Stromal cell/ T cell/ pan-APC | 2851 |
| 8899_AS_6 | Vanderbilt HTAN | Tumor cell/ Normal epithelial cell/ Stromal cell/ T cell | 2669 |
| 8899_AS_7 | Vanderbilt HTAN | Normal epithelial cell/ Stromal cell/ T cell/ pan-APC | 3182 |
| SH-16-04266-A1 | Fred Hutch | T cell/ pan-APC | 2607 |
| SH-16-07447 | Fred Hutch | Tumor cell/ T cell | 3544 |
| SH-17-06079-B1 | Fred Hutch | Tumor cell/ T cell/ pan-APC | 922 |
| SH-17-06138-A1 | Fred Hutch | T cell | 2427 |
| SU-15-18753-A1 | Fred Hutch | T cell | 1961 |
| SU-15-19531-A1 | Fred Hutch | Tumor cell/ T cell/ pan-APC | 1818 |
| SU-15-27301-B1 | Fred Hutch | Normal epithelial cell/ T cell | 66 |
| SU-16-02468-B1 | Fred Hutch | T cell | 151 |
| SU-17-05594-A1 | Fred Hutch | Tumor cell/ T cell/ pan-APC | 688 |
| SU-17-07002-A1 | Fred Hutch | T cell/ pan-APC | 366 |
| SU-17-09232-A1 | Fred Hutch | T cell | 1434 |
| SU-17-14212-A1 | Fred Hutch | T cell/ pan-APC | 1890 |
| SU-17-15020-C1 | Fred Hutch | Tumor cell/ T cell/ pan-APC | 384 |
| SU-17-18554-B1 | Fred Hutch | Tumor cell/ T cell | 2333 |
| SU-17-23223-A1 | Fred Hutch | Tumor cell/ Stromal cell/ T cell/ pan-APC | 776 |
| SU-17-25332-D1 | Fred Hutch | Tumor cell/ T cell/ pan-APC | 798 |
| SU-17-25385-B1 | Fred Hutch | Tumor cell/ T cell | 2816 |
| SU-17-30257-A1 | Fred Hutch | Tumor cell/ Stromal cell/ T cell/ pan-APC | 1107 |
| TENX152 | HEST-1K | Tumor cell/ Normal Epithelial cell/ Stromal cell/ T cell/ pan-APC | 4096 |
| TENX28 | HEST-1K | Tumor cell/ Normal Epithelial cell/ T cell/ pan-APC | 3134 |
| TENX29 | HEST-1K | Tumor cell/ pan-APC | 3134 |
| TENX49 | HEST-1K | Tumor cell/ Normal Epithelial cell/ Stromal cell/ T cell/ pan-APC | 2657 |
| TENX89 | HEST-1K | Normal Epithelial cell | 5938 |
| TENX90 | HEST-1K | Normal Epithelial cell | 6042 |

**Table S1 continued**

| Sample ID | Source | Training for | Total Tiles |
|---|---|---|---|
| ZEN38 | HEST-1K | Tumor cell/ Stromal cell/ T cell/ pan-APC | 387 |
| ZEN39 | HEST-1K | Stromal cell | 1219 |
| ZEN42 | HEST-1K | Tumor cell/ Stromal cell/ T cell | 1192 |
| ZEN43 | HEST-1K | Tumor cell/ Stromal cell/ T cell/ pan-APC | 691 |
| ZEN44 | HEST-1K | Tumor cell/ Normal Epithelial cell/ Stromal cell/ pan-APC | 1048 |
| ZEN45 | HEST-1K | Tumor cell/ Normal Epithelial cell/ Stromal cell/ pan-APC | 328 |
| ZEN46 | HEST-1K | T cell/ pan-APC | 1803 |

#### Table S2. Training samples and tile counts (Breast cancer).

| Sample ID | Source | Training for | Total Tiles |
|---|---|---|---|
| 06092021 | 10x Genomics | Tumor cell/ Stromal cell/ pan-APC/ T cell/ Normal Epithelial cell | 2428 |
| 06232020-S1 | 10x Genomics | Stromal cell/ pan-APC/ T cell/ Normal Epithelial cell | 3683 |
| 06232020-S2 | 10x Genomics | Stromal cell/ pan-APC/ T cell/ Normal Epithelial cell | 3866 |
| 07012022 | 10x Genomics | pan-APC/ T cell/ Normal Epithelial cell | 4000 |
| 10272020 | 10x Genomics | Stromal cell/ Normal Epithelial cell | 3860 |
| GSM5732357_A | Liu et al. | pan-APC/ T cell/ Normal Epithelial cell | 2859 |
| GSM5732358_B | Liu et al. | Tumor cell/ Stromal cell/ pan-APC/ T cell | 4222 |
| GSM5732359_C | Liu et al. | Tumor cell/ Stromal cell/ pan-APC/ T cell | 4615 |
| GSM5732360_D | Liu et al. | Tumor cell/ pan-APC/ T cell/ Normal Epithelial cell | 4049 |
| 1142243F | Wu et al. | pan-APC/ T cell | 4517 |
| 1160920F | Wu et al. | pan-APC/ T cell/ Normal Epithelial cell | 4655 |

**Table S3. Cox proportional hazards regression models of median distance features for progression-free survival in TCGA breast cancer.** Hazard ratios with 95% confidence intervals and $p$-values are reported from the Cox proportional hazards model (either unadjusted or adjusted) with progression-free survival as the outcome. Significant associations ($p < 0.05$) are shown in bold.

| | Unadjusted Cox Model | | Adjusted Cox Model | |
|---|---|---|---|---|
| Median Distance | HR (95% CI) | $p$-value | HR (95% CI) | $p$-value |
| Tumor cells to Stromal cells | 0.912 (0.440–1.890) | 0.805 | 0.733 (0.314–1.711) | 0.473 |
| Tumor cells to Pan-APCs | 0.956 (0.599–1.524) | 0.849 | 0.873 (0.508–1.500) | 0.624 |
| Tumor cell to T cells | 0.845 (0.346–2.063) | 0.712 | 0.320 (0.116–0.886) | **0.028** |
| Stromal cells to Tumor cells | 0.914 (0.753–1.108) | 0.358 | 1.066 (0.878–1.294) | 0.517 |
| Stromal cells to pan-APCs | 1.301 (0.695–2.433) | 0.411 | 1.016 (0.493–2.092) | 0.966 |
| Stromal cells to T cells | 0.563 (0.274–1.156) | 0.118 | 0.442 (0.193–1.015) | 0.054 |
| pan-APCs to Tumor cells | 0.845 (0.669–1.067) | 0.156 | 0.994 (0.798–1.238) | 0.958 |
| pan-APCs to Stromal cells | 1.303 (0.832–2.043) | 0.248 | 1.083 (0.554–2.117) | 0.815 |
| pan-APCs to T cells | 0.403 (0.170–0.956) | **0.039** | 0.434 (0.177–1.068) | 0.069 |
| T cells to Tumor cells | 0.934 (0.739–1.179) | 0.565 | 1.008 (0.814–1.249) | 0.942 |
| T cells to Stromal cells | 1.227 (0.503–2.998) | 0.653 | 0.866 (0.302–2.486) | 0.790 |
| T cells to pan-APCs | 0.868 (0.388–1.940) | 0.730 | 0.841 (0.365–1.939) | 0.685 |
